# Identification of RNA binding proteins that mediate a quality control mechanism of splicing

**DOI:** 10.1101/2025.07.20.665773

**Authors:** Maram Arafat, Valer Gotea, Mubarak I Umar, Aya Muharram, Eyal Kamhi, Reuven Agami, Joseph Sperling, Markus Hafner, Laura Elnitski, Ruth Sperling

## Abstract

Accurate splicing, which involves the controlled removal of non-coding sequences (introns) from precursor messenger RNAs (pre-mRNAs), is essential for producing correct mature mRNAs that encode functional proteins. Within pre-mRNAs, latent splice sites (LSSs) resemble proper splice sites but are usually not used because their activation can introduce in-frame STOP codons. The nuclear suppression of splicing (SOS) mechanism prevents the use of LSSs. Although the SOS mechanism is not fully understood, recent studies have identified initiator-tRNA and the NCL protein as key components. To discover additional regulators, we performed a genetic screen targeting RNA-binding proteins (RBPs) with an siRNA library and a luminescence reporter for latent splice site activation. This identified five RBPs — ALYREF (THOC4), PPIE, DDX41, DHX38, and HNRNPA2B1 — whose knockdown significantly increased LSS usage in the reporter. RNA-Seq analysis after knocking down each of these RBPs confirmed these results, showing widespread LSS activation in hundreds of mRNAs. Among these, we focused on ALYREF, a conserved protein involved in mRNA export and splicing. Using fPAR-CLIP, we found that U5 snRNA is ALYREF’s main binding partner. Overexpressing ALYREF deletion mutants activated latent splicing, and affinity purification confirmed its interaction with U5 snRNA. These mutants exhibited different binding properties, highlighting the importance of specific structural elements within ALYREF in SOS regulation. Our findings reveal that nuclear RBPs play a key role in suppressing LSS activation and suggest that ALYREF has a novel role in maintaining splicing accuracy within the spliceosome, advancing our understanding of the SOS mechanism.

## Introduction

In higher eukaryotes, the generation of correct mature messenger RNA (mRNA) transcripts depends on the precise splicing of precursor mRNA (pre-mRNA). This process involves removing non-coding intronic sequences and assembling exons into the final mRNA product (Berget et al. 1977; Chow et al. 1977; Darnell 1978). Alternative splicing (AS) significantly increases mRNA and protein diversity by allowing for the selective inclusion or exclusion of exons. Despite its biological importance, many aspects of this complex and highly regulated process are still not well understood. A key step in this process is choosing the donor and acceptor splice sites (i.e., 5’SS and 3’SS), which define the boundaries of the intron to be removed. While the mechanism of single-intron transcript splicing has been extensively characterized (Konarska et al. 1985; Papasaikas and Valcarcel 2016; Wachutka et al. 2019), many aspects of the regulation of AS still require elucidation (Sperling 2017). Understanding these aspects will help explain how changes in AS influence processes such as apoptosis and cell proliferation, and how they contribute to numerous human diseases, including cancer (Kelemen et al. 2013; Lee and Rio 2015; Chabot and Shkreta 2016; Papasaikas and Valcarcel 2016).

Accumulating evidence suggests the presence of a nuclear mechanism known as suppression of splicing (SOS), which regulates splice site selection to prevent the utilization of intronic splice sites that could introduce premature in-frame stop codons. This mechanism operates independently of nonsense-mediated decay (NMD) and other degradation pathways (Li et al. 2002; Miriami et al. 2002; Wachtel et al. 2004; Kamhi et al. 2010; Nevo et al. 2012; Arafat and Sperling 2022), and is confirmed by studies in nematode mutant strains lacking essential NMD genes (Nevo et al. 2015). The significance of the SOS mechanism is highlighted by the presence of millions of 5’SS-like sequences within intronic regions that remain unused under normal growth conditions (latent splice sites, LSS) (Li et al. 2002; Miriami et al. 2002; Kamhi et al. 2006; Sperling and Sperling 2008; Kamhi et al. 2010; Nevo et al. 2012). Use of the overwhelming majority (>98%) of LSSs as 5’SS would result in the incorporation of in-frame STOP codons into the mRNA, a deleterious process observed under cellular stress and in cancer (Nevo et al. 2012). Although the exact mechanisms of SOS are not yet fully understood, critical roles have been identified for initiator-tRNA (ini-tRNA), which, apart from its cytoplasmic function in protein translation, is involved in base pairing with the AUG sequence in the nucleus (Kamhi et al. 2010). This interaction, likely occurring in a complex with auxiliary proteins, establishes a register for reading frame recognition by SOS (Sperling and Sperling 2008; Kamhi et al. 2010; Sperling 2019). Recently, nucleolin (NCL) was identified as the first protein component associated with SOS (Shefer et al. 2022) through: (*i*) NCL’s direct and specific interaction with ini-tRNA in the nucleus, but not in the cytoplasm; (*ii*) its association with ini-tRNA and pre-mRNA in spliceosomal fractions; (*iii*) demonstrating that the recovery of suppression of latent splicing by ini-tRNA complementation is NCL dependent; and (*iv*) activation of hundreds of latent splice sites in coding transcripts upon NCL knockdown. We suspect that NCL operates together with other molecular components that still need to be identified to gain a complete understanding of the SOS mechanism.

In this study, we searched for protein components of the SOS mechanism by developing a luminescence-based reporter system, which monitors LSS activation. Using this reporter system, we screened a library of siRNAs targeting RNA-binding proteins (RBPs) to identify key players in this process. The screen identified five prominent candidates: ALYREF (also known as THOC4), PPIE, DDX41, DHX38, and hnRNPA2B1. All five candidate proteins are nuclear and have previously been implicated in splicing, either through direct interactions with the spliceosome or through functional roles in pre-mRNA processing. The evolutionarily conserved ALYREF is an abundant RBP and part of the TREX complex required for mRNA export. ALYREF is recruited to mRNA during splicing and becomes a core component of the exon junction complex (EJC) (Masuda et al. 2005; Chi et al. 2013). Peptidylprolyl isomerase E (PPIE), a member of the peptidyl-prolyl cis-trans isomerase (PPIase) family, contains an RNA-binding domain and was previously implicated in splicing (Thapar 2015). DEAD-box helicase 41 (DDX41) has been linked to the spliceosome through mass spectrometry analyses, and its mutations have been associated with myelodysplastic syndromes that are characterized by splicing factor mutations (Polprasert et al. 2015). DEAH-box helicase 38 (DHX38), an ATPase member of the DEAD/H box family of splicing factors, is essential for the catalytic step II in the pre-mRNA splicing process (Zhou and Reed 1998). It appears to interact transiently with the spliceosome through its non-conserved N-terminal domain (Schwer and Guthrie 1991; Wang and Guthrie 1998; Wang et al. 1998). Finally, heterogeneous nuclear ribonucleoprotein A2/B1 (HNRNPA2B1), part of the A/B subfamily of ubiquitously expressed heterogeneous nuclear ribonucleoproteins, associates with pre-mRNAs in the nucleus and plays a significant role in pre-mRNA processing, metabolism, and transport (Han et al. 2010). We further demonstrate by RNA-seq that the knockdown of each of these five splicing-associated factors results in the activation of hundreds of LSSs, introducing in-frame STOP codons and implicating these proteins in splice-site selection

We focused on studying ALYREF, due to its known interactions with the spliceosome and the export machinery (Masuda et al. 2005; Chi et al. 2013). We comprehensively identified ALYREF binding sites on its RNA targets with nucleotide resolution using fPAR-CLIP [fluorescent photoactivatable ribonucleoside enhanced crosslinking and immunoprecipitation (Anastasakis et al. 2021)], revealing U5 snRNA as the primary partner of ALYREF. Affinity purification of nuclear complexes assembled with overexpressed wild-type (WT) ALYREF confirmed its binding to U5 snRNA. Furthermore, overexpression of ALYREF deletion mutants resulted in the activation of latent splicing, similar to the effects observed with ALYREF knockdown. ALYREF deletion mutants also showed differences in their binding to U5 snRNA and other spliceosome components, indicating the absence of elements essential for SOS regulation. In addition to nominating new components of the SOS, our study demonstrates a novel role for ALYREF in splice site selection and SOS through its binding to U5 snRNA, an important component of the spliceosome throughout all stages of the splicing reaction. More broadly, these results support a model in which networks of abundant nuclear RBPs suppress latent splicing events. The diversity of these RBPs likely ensures that the SOS mechanism operates robustly across variable sequence contexts, safeguarding transcriptome fidelity through cooperative surveillance.

## Results

### A reporter-based RNAi screen identifies RNA-binding proteins involved in SOS

RNA binding proteins (RBPs) are crucial regulators of AS (Kamhi et al. 2010; Kelemen et al. 2013; Akerman et al. 2015; Papasaikas and Valcarcel 2016; Fiszbein and Kornblihtt 2017; Sperling 2017; Dvinge 2018), and we reasoned that they likely also play a role in mediating the SOS (Kamhi et al. 2010). To search for RBPs involved in SOS, we used a previously established siRNA library specifically focused on RBPs (Jenal et al. 2012) and used it with a newly developed luminescence-based reporter system, named IRS1 (IRES-based Reporter for SOS #1) (**Figure 1A**). This system is based on a minigene derived from the CAD gene (Carbamoyl-Phosphate Synthetase 2, Aspartate Transcarbamylase, and Dihydroorotase). This minigene contains a latent 5’ splice site (LSS) within its second intron, which is suppressed by SOS (Li et al. 2002; Wachtel et al. 2004; Kamhi et al. 2006; Kamhi et al. 2010). We cloned a Renilla luciferase gene downstream of the intron, directing its translation via the internal ribosome entry site (IRES) of the cricket paralysis virus (CrPV) (Petersen et al. 2006). The design allows the Renilla luciferase sequence to be split between the 3’ end of the latent exon (residues 1-40; **Figure 1A**, bold green line) and the 5’ end of the downstream exon (residues 41-190; **Figure 1A**, bold blue line). Note that disrupting the pseudoknot (PK) structure in this region inhibits IRES-mediated translation (Jan and Sarnow 2002; Schuler et al. 2006). The open reading frame (ORF) of the Renilla luciferase gene is positioned out of frame with the initial ATG start codon. Consequently, during normal splicing, Renilla expression remains minimal. Only latent splicing produces an intact CrPV IRES (residues 1-190; **Figure 1A**, magenta heavy line), resulting in Renilla luciferase expression. Thus, latent splicing levels can then be quantified by quantifying Renilla luciferase activity and normalizing it to the activity of Firefly luciferase, which is expressed from an independent promoter and not influenced by Renilla luciferase levels. We generated three additional control constructs (**Figure 1B**). The two positive controls, IRS2 and IRS3, carry point mutations that change the translation start codon from ATG to ACG and AAA, respectively, and activate latent splicing (Kamhi et al. 2006; Kamhi et al. 2010). The third mutant, IRS6, served as a negative control and carries a CAG to GCA mutation at the 3’SS used for both authentic and latent splicing, thus inactivating both splicing events.

**Figure 1.**
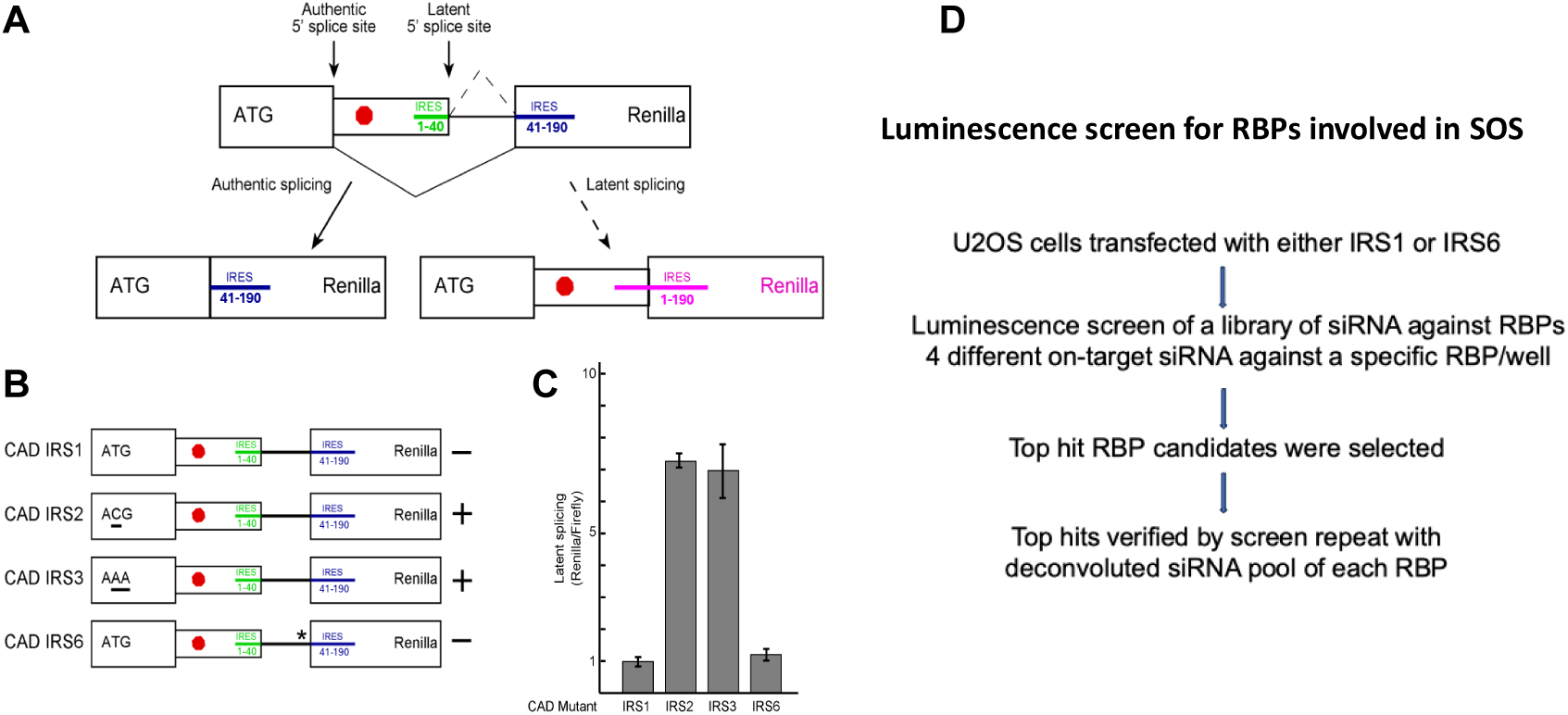
Renilla expression from the IRES reporter system. **(A)** A schematic representation of the IRS reporter. The scheme depicts the cloned pre-mRNA where the sequence of the CrPV IRES is split between the latent exon (green) and the downstream exon (blue). The two possible 5’ alternative splicing events are illustrated. When splicing occurs at the authentic 5’ splice site (left), no Renilla is expressed because its sequence is out of frame of the first AUG, and it cannot be expressed from the CrPV IRES as the transcript lacks the essential nucleotides from the 5’ end of that IRES. However, when latent splicing occurs (right), Renilla is expressed through an intact CrPV IRES (magenta). Large box-exon; small box-latent exon; colored lines - indicated split IRES sequences; ATG – initiation codon sequence; red octagon – in-frame stop codon. (**B**) Schematic drawings of the four Renilla constructs: IRS1, the reporter construct; IRS2 and IRS3, constructs in which the start codon sequence was mutated from ATG to ACG, or AAA, respectively; IRS6, in which the 3’ splice site was mutated. Expected Renilla expression (+) or lack of expression (-) from the IRS constructs is indicated on the right. **(C)** HEK 293T cells were transfected with each of the indicated plasmids. Forty-eight hr. post-transfection luminescence levels were measured from Renilla and Firefly luciferase (Promega Dual-Luciferase system). Levels of latent splicing are expressed as the ratio between the luminescence of Renilla luciferase to that from the Firefly luciferase. As expected, the Renilla luciferase levels increase significantly (p<0.01) in cells transfected with IRS constructs having a mutation in the start codon (ATG->ACG, ATG->AAA; IRS2 and IRS3 respectively), as a consequence of latent splicing. Background levels are observed in the cases of IRS1 and IRS6. (**D**) Scheme of the luminescence screen for RBPs involved in SOS.

The four IRS constructs were individually transfected into HEK 293T cells, and 48 h post-transfection, the Renilla and Firefly luciferase output was measured. As expected, mutation of the start codon (both IRS2 and IRS3) resulted in a substantial increase in Renilla luciferase expression. As expected, a low Renilla signal was obtained with IRS1 under normal growth conditions, indicating minimal latent splicing, also seen in the negative control IRS6. Under heat shock, which suppresses SOS and activates latent splicing (Miriami et al. 1994; Nevo et al. 2012; Nevo et al. 2015), IRS1 showed a robust Renilla signal (Supplementary Figure S1), demonstrating its utility as a reporter for screening activators of latent splicing.

Using our reporter system, we calibrated and conducted a high-throughput genetic screen in human osteosarcoma (U2OS) cells to identify RBPs that suppress SOS and activate latent splicing (see Materials and Methods, and **Figure 1D**). Cells grown in 384-well plates were transiently transfected with either the IRS1 or IRS6 plasmids, and RBPs were knocked down using a pool of four different siRNAs. After seventy-two hours, the cells were harvested, and we measured the Renilla to Firefly luciferase ratio. We calculated the average activation of latent splicing following RBP knockdowns by normalizing the luciferase signal from the IRS1 sample against the negative control (IRS6). Positive hits were identified only if the luciferase increase was observed in both replicates and was at least one standard deviation greater than the average signal from the entire microplate. This screening revealed ten RBPs whose knockdown led to the activation of latent splicing (see **Table 1** and Supplementary Table S1).

**Table 1.** Top hits of RBP screen and results of deconvolution validation.

| No | Protein name | IRS1/IRS6 Ave | Remarks | Deconvolution |
| --- | --- | --- | --- | --- |
| 1 | RBP2 | 1.66 | Intracellular transport of retinol | 2/4 |
| 2 | **DHX38** | **1.52** | **Prp16 helicase** | **2/4** |
| 3 | **THOC4** | **1.48** | **ALY/REF export and splicing protein** | **4/4** |
| 4 | RBM27 | 1.47 | RNA binding protein | 2/4 |
| 5 | **PPIE** | **1.46** | **Catalyzes isomerization of prolines** | **4/4** |
| 6 | DHX37 | 1.43 | DEAD box RNA helicase | 1/4 |
| 7 | RBMY1B | 1.4 | RNA Binding Motif Protein Y-Linked Family 1 member B, with SRGY boxes in C-terminus | 0/4 |
| 8 | BAMBI | 1.38 | BMP And Activin Membrane Bound Inhibitor | 1/4 |
| 9 | **DDX41** | **1.34** | **DEAD box helicase** | **2/4** |
| 10 | CBX8 | 1.32 | Component of a gene repressor complex | 1/4 |

We validated the results of our high-throughput screening by measuring latent splice site activation in our reporters after knockdown of our ten identified hits with each individual siRNA from the initial pool of four. For validation, we required that at least two siRNAs activate latent splicing. This rescreen revealed six hits: RBP2, DHX38, THOC4, RBM27, PPIE, and DDX41 (**Table 1**). Knockdown of negative control RBPs, UPF1 (RENT1) and ELAVL3, did not activate latent splicing, confirming our previous results that SOS is indeed not dependent on UPF1 and thus NMD (Wachtel et al. 2004; Nevo et al. 2015).

In addition to the six hits from the screen, we also tested the effect of three heterogeneous nuclear ribonucleoproteins (hnRNPs) - HNRNPA1, HNRNPA2B1, and HNRNPC - on our reporter system as well as on the CAD minigene (Li et al. 2002), considering their well-established roles in splicing regulation and nuclear mRNA processing. Of these additional targets, only HNRNPA2B1 knockdown resulted in latent splice site activation (**Figure 2**), further highlighting the variability in function of the hnRNPs (Geuens et al. 2016). RT-PCR after knockdown of DHX38 confirmed the expected increase in reporter mRNA levels (**Figure 2E**). Because of their well-established role in splicing (Schwer and Guthrie 1991; Masuda et al. 2005; Chi et al. 2013; Polprasert et al. 2015; Thapar 2015), we concentrated our follow-up efforts on characterizing ALYREF, PPIE, DHX38, DDX41, and HNRNPA2B1 as potential components of the SOS system.

**Figure 2.**
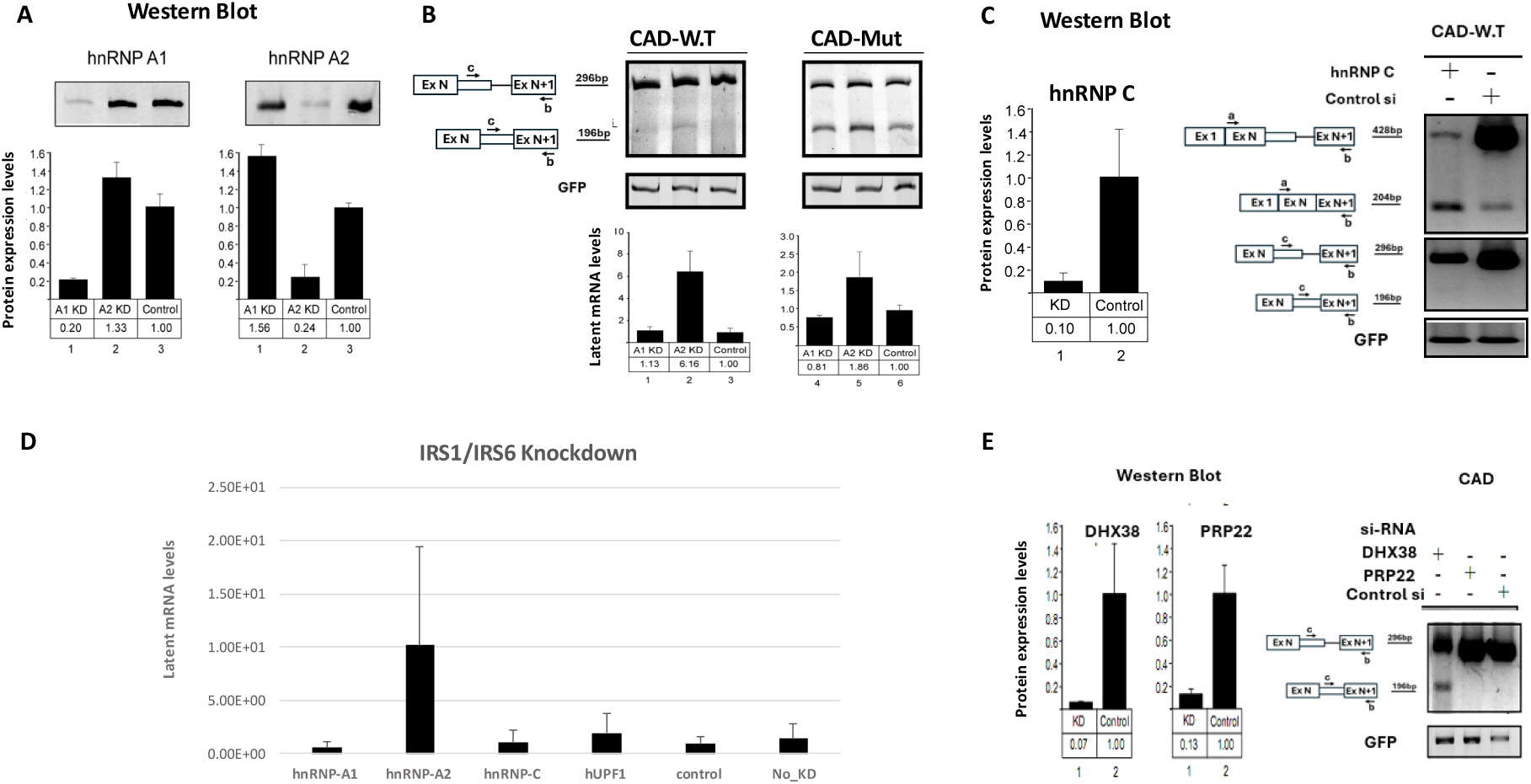
Knockdown of DHX38 and hnRNPA2B1 activates latent splicing. HEK293 cells were treated with the relevant siRNA. Knockdown was analyzed by WB, and latent splicing activation of CAD minigene transcripts were analyzed by RT-PCR. **(A, B)** KD of hnRNPA2B1(hnRNP A2), but not hnRNPA1 activates latent splicing. (**A**) WB, (**B**) RT-PCR. A positive control of CAD Mut (CAD mutant lacking all stop codons) in which latent splicing is activated, shown for comparison. (**C**) KD of hnRNP C does not activate latent splicing. (left) WB results; (right) RT-PCR results. (**D**) KD of hnRNPA2B1 activates latent splicing using the IRS1/IRS6 system (see text). (**E**) KD of DHX38 but not Prp22 activates latent splicing. (left) WB results; (right) RT-PCR.

### Analyzing the impact of knockdown of potential SOS factors on latent splicing

We aimed to investigate the effectiveness of our five candidate genes in globally suppressing latent splicing. Therefore, we conducted RNA-Seq analysis following siRNA knockdown in HeLa cells. The changes in gene expression for each mRNA were calculated using DESeq2, confirming that the knocked-down five factors were expressed well below the controls (with one exception) (**Figure 3**). To identify activated LSSs in the RNA-seq reads, we used the QoRTs tool (Hartley and Mullikin 2015) to quantify split reads, i.e., sequence reads spanning exon junctions (**Table 2**).

**Figure 3.**
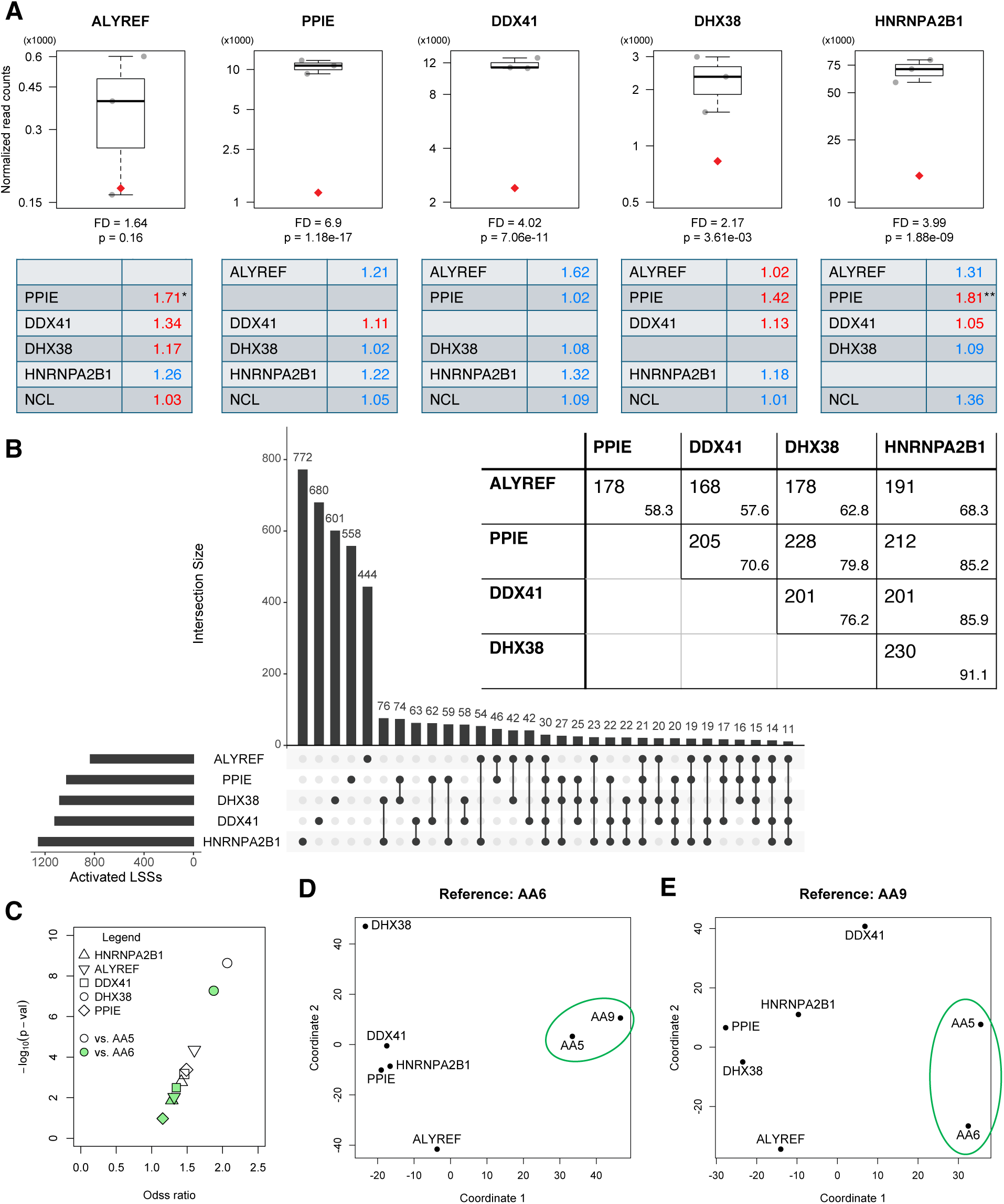
Outcome of siRNA knockdown of five splicing factors. (**A**) Gene expression levels of the transcripts of the six factors upon specific siRNA knockdown (red diamond) compared to AA4-AA6 control samples (gray circles). The targeted factor in each experiment is indicated above the graph, whereas decrease in level for that factor is indicated below the graph (FD: fold difference as estimated through DESeq2 analysis; associated p-value un-adjusted for multiple testing is indicated below the FD value). Changes in expression level for the other five (non-targeted) factors for each experiment are indicated in the corresponding table below the graph. Blue – upregulation, red – downregulation. Significant changes in expression levels are indicated by * (* - 0.05; ** - 0.01 significance level). (**B**) UpSet plot with the shared LSSs activated relative to the AA5 control upon specific factor knockdown. Pairwise figures are shown in the insert table: observed: upper left; expected: lower right (see Methods). (**C**) Volcano-style plot of magnitude of effect (odds ratio) and significance (p-value computed with one-sided Fisher’s exact test) of activated LSSs compared to adSS_3n_ in samples with siRNA targeted factors relative to AA5 and AA6 control samples. (**D, E**) Multidimensional scaling using Jaccard distances based on shared activated LSSs relative to AA6 and AA9 control samples, respectively.

**Table 2.** Summary of RNA-seq reads and split read support for LSSs.

| Sample type | Sample name | Uniquely mapped reads | Number of LSSs supported by split reads | Number of LSSs supported by 4 or more split reads |
| --- | --- | --- | --- | --- |
| Targeted siRNA | ALYREF | 105,888,093 | 21,506 | 5,889 |
|  | PPIE | 97,842,700 | 20,507 | 5,575 |
|  | DDX41 | 125,892,498 | 29,295 | 6,925 |
|  | DHX38 | 97,007,698 | 17,743 | 5,206 |
|  | HNRNPA2B1 | 122,307,030 | 23,536 | 6,485 |
| Non-specific siRNA | AA4 | 124,617,677 | 20,362 | 5,859 |
|  | AA5 | 103,617,575 | 23,708 | 5,889 |
|  | AA6 | 100,944,977 | 18,473 | 5,516 |
| Mock transfection | AA9 | 101,073,720 | 18,562 | 5,234 |

Across all samples, we identified over 94,000 unique active LSSs, each supported by at least one split read. This represents nearly 7% of our total of 1.34 million curated LSSs, and is consistent with our previous report that identified 70,000 LSSs upon NCL knockdown (Shefer et al. 2022). When we increased the stringency of our criteria to require at least four split reads at every LSS, we still found over 5,000 LSSs activated by the knockdown of any of our five RBPs. Notably, LSSs supported by at least four split reads exhibited higher overlap rates than those supported by a single read (Jaccard index of ∼24% vs. ∼20%, respectively), consistent with higher probabilities of capturing transcripts expressed at higher levels. To visualize the global similarity of splicing changes, we performed multidimensional scaling on shared LSS activation profiles and excluded the control sample affecting NCL, as NCL is itself a core regulator (**Figure 3C-E**).

If our candidate RBPs have a role in SOS, their knockdown should result in significantly elevated levels of activated LSSs. To test this prediction, we followed a computational approach we developed previously (Shefer et al. 2022) and calculated the ratio of activated LSS to activated alternative donor splice sites (adSS_3n_), which maintain the open reading frame and lack upstream in-frame STOP codons and thus should not elicit SOS. We found that the knockdown of our RBP candidates resulted in an approximately two-fold increase in the ratio of activated LSSs to activated adSS_3n_ events, compared to controls (**Figure 3B**). These differences were significant (one-sided Fisher’s exact test), with the exception of the PPIE knockdown (significant vs. one control only), consistent with the involvement of all five factors in the SOS mechanism.

Next, we investigated commonalities between LSSs found to be activated in multiple samples. The proportion of overlap between activated LSSs in this stringent, normalized pipeline is smaller than what we observed for LSSs supported by at least four reads (a less stringent approach), with Jaccard similarity indices between 9.3% (DDX41 & HNRNPA2B1) and 12.1% (DHX38 & PPIE). However, the number of common activated LSSs between samples is severalfold (2.3–3.1) higher than what can be expected based on random sampling of all possible LSSs (see Methods, **Figure 3B**), suggesting a shared functional splicing mechanism was affected by the knockdown of the five targeted factors. Moreover, multidimensional scaling of shared patterns among activated LSSs (see Methods) reveals that control samples cluster separately from case samples, regardless of whether the reference sample was transfected with a non-specific siRNA or mock-transfected (**Figure 3**). This observation indicates that specific targeting of factors results in non-random LSS activation patterns, consistent with a common underlying mechanism involving these factors (Scacheri et al. 2004).

### Independent evaluation of splicing activation through RT-PCR

We validated the activation of specific LSSs identified through RNA sequencing after the knockdown of our SOS candidate RBPs (see **Figure 4A** for an example). We knocked down ALYREF, PPIE, hnRNPA2B1, and DHX38 in HEK293, and additionally targeted ALYREF and hnRNPA2B1 in HeLa cells (**Figure 4B**) to ensure that our results were cell-type independent. We then conducted RT-PCR experiments to examine a subset of activated LSSs across three biological replicates. For each SOS factor, we selected at least four transcripts that showed the strongest evidence of LSS activation according to the RNA-seq data (**Figure 4C,D**). In each case, we observed a significant increase in LSS usage (the identity of the bands was verified by sequencing).

**Figure 4.**
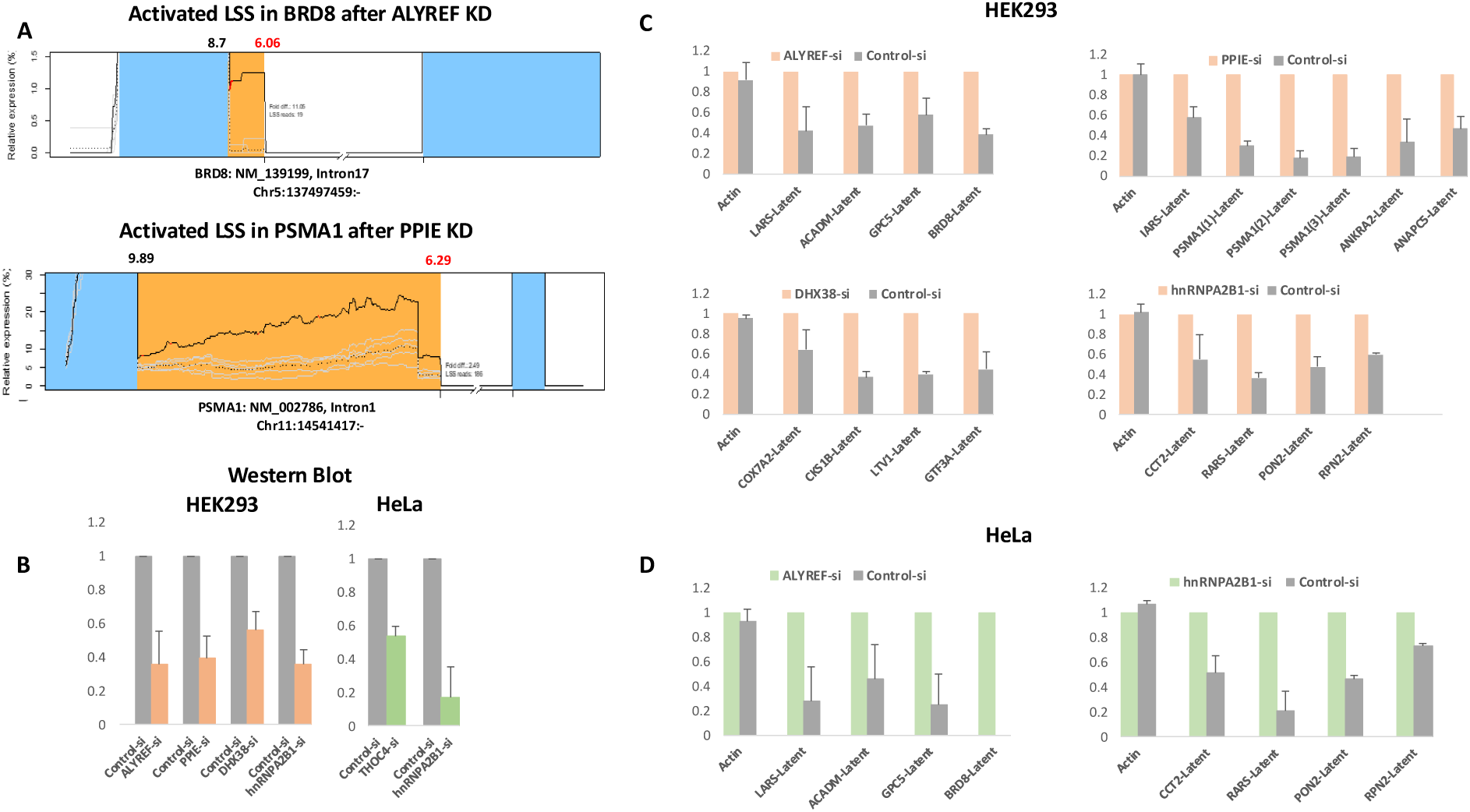
Activation of latent splicing at LSSs in top gene targets expressed in cells in which ALYREF, or PPIE, or DHX38, or hnRNPA2B1 were knocked down. RT-PCR validation of activation of latent splicing at LSSs in top gene targets following factor knock down. CONTsi treated cells were used as control. (**A**) Expression profiles of exon extension regions upstream of LSSs activated in BRD8 after ALYREFsi (top), and PSMA1 after PPIEsi (Bottom). Gray lines correspond to levels observed in control samples, and the dashed lines corresponds to the average values in control samples. Positions of in-frame STOPs are shown by red segments. Numbers on top of the graph indicate MaxEntScan scores for the annotated 5’SS (black) and the LSS (red). The length of the exon extension is indicated in the corresponding segment of the gene model. (**B**) Western blot analysis of knockdown of each of the four gene transcripts. (**C**) RT-PCR validation in HEK293 cells of LSSs activation in top targets after knockdown of each of the four gene transcripts. Bars represent averages and SEMs for three biological replicates. (**D**) RT-PCR validation in HeLa cells of LSSs activation in top targets after knockdown of either ALYREF or hnRNPA2B1. Bars represent averages and SEMs for three biological replicates.

### Spliceosomal RNA targets and binding sites of ALYREF-fPAR-CLIP analysis

We further explored the role of ALYREF in SOS in more depth. ALYREF was one of our top hits on the screen and was previously shown to be part of the supraspliceosome, the structure required for SOS (Kotzer-Nevo et al. 2014). Affinity purification of supraspliceosomes (Spann et al. 1989; Azubel et al. 2006) assembled on a PP7-tagged SMN1 transcript expressed from an SMN1 minigene stably expressed from HeLa cells revealed that ALYREF is associated with the affinity-purified SMN1 supraspliceosomes together with splicing factors (Supplementary **Figure S2**).

To identify ALYREF binding sites on mRNAs at nucleotide resolution across the transcriptome, we used fPAR-CLIP (Anastasakis et al. 2021). This technique relies on photocrosslinking of RNAs metabolically labeled with 4-thiouridine (4SU) with interacting RBPs by UV light (l > 320 nm). To specifically study ALYREF binding sites within the supraspliceosome, we isolated nuclear supernatants (Spann et al. 1989; Azubel et al. 2006), immunoprecipitated ALYREF, and ligated fluorescent adapters to the crosslinked RNA. This was followed by fractionation on a denaturing polyacrylamide gel (see **Figure 5A**). As anticipated, we observed a fluorescent band at 47 kDa, which corresponds to the expected molecular weight of the adapter-ligated ALYREF-RNP in our sample, but not in the control from non-crosslinked cells (**Figure 5A,B**). We then isolated RNA and performed deep sequencing from two biological replicates. The sequenced reads were aligned to the human genome and grouped into clusters to identify those enriched for crosslink-induced T-to-C mutations (Hafner et al. 2010; Anastasakis et al. 2021) (Supplementary **Figure S3)**. The two biological replicates demonstrated excellent concordance, with a Spearman correlation coefficient of 0.76 (**Figure 5C**). Overall, we identified >20,000 clusters that predominantly mapped to pre-mRNAs (**Figure 5D**), along with exonic sequences and non-coding RNAs (Supplementary Table S2).

**Figure 5.**
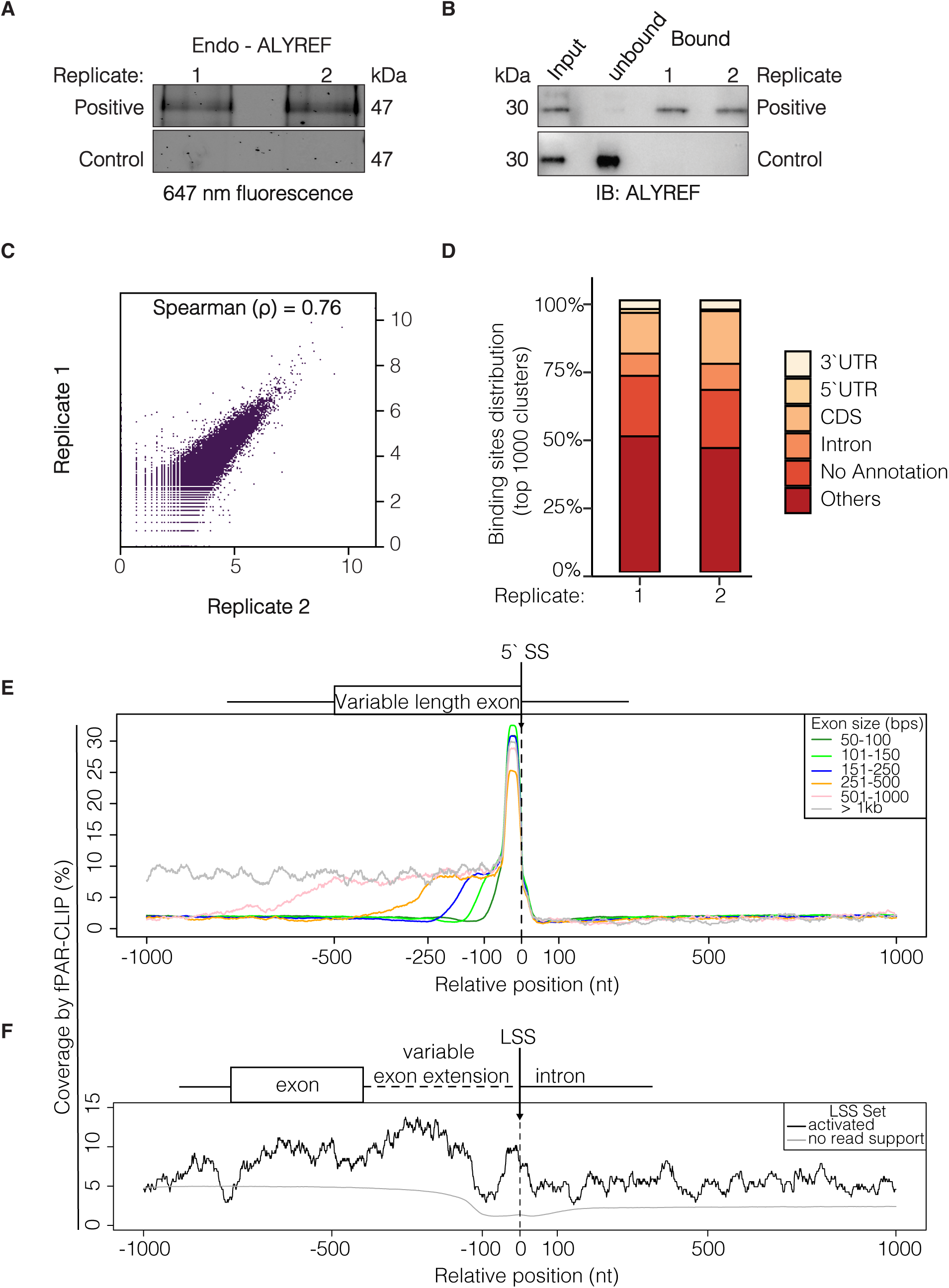
ALYREF RNA binding analysis. (**A**) Fluorescent image showing crosslinked, ribonuclease treated, and fluorescent adapter ligated endogenous ribonucleoproteins (RNPs) after ALYREF IP from HEK293 cells, separated by SDS-PAGE. (**B**) Immunoblot control demonstrating successful immunoprecipitation of endogenous ALYREF in positive samples and not in control samples. (**C**) Spearman correlation coefficient (ρ) analysis of two biological replicates of ALYREF fPAR-CLIP, showing a strong reproducibility with a ρ of 0.76. (**D**) Distribution of endogenous ALYREF binding sites across various RNA annotations. Details of the genes within each annotation are provided in Supplementary **Table S2**. (**E,F**) fPAR-CLIP profiles of genomic regions in HEK 293 cells. (**E**) fPAR-CLIP profiles of internal exons and downstream intronic regions. (**F**) fPAR-CLIP profiles centered on LSSs that are activated (i.e., are spliced in RNA-seq data) upon knockdown of THOC4/ALYREF in HeLa cells (305), and of potential LSSs that exhibit no split read support in the RNA-seq data in HeLa cells (942,857). To minimize the influence of upstream exonic regions, these sets only include LSSs that are located at least 100 nt downstream from the annotated 5’SS.

Metagene analysis revealed that ALYREF preferentially bound just upstream of the 5’SS and, to a lesser degree, across the exon (**Figure 5E**), consistent with previous results showing its binding to the exon junction (Viphakone et al. 2019). We then focused specifically on LSSs activated upon ALYREF knockdown. Our results showed weaker binding upstream of the LSS, along with an elevated signal also upstream of the LSS, similar to what we observed for the 5’SSs (**Figure 5F**). In contrast, LSSs without split-read support exhibited a significantly weaker signal. Consistent with an intrinsic interaction of ALYREF with the spliceosome, the top RNA target was U5 snRNA (RNU5-D1) (**Figure 6**), bound specifically at the base of a stem (at U73) (**Figure 6B,C** and Supplementary **Figure S3C, Table S2**). Consistent with our previous findings of ini-tRNA involvement in SOS (Li et al. 2002; Wachtel et al. 2004; Kamhi et al. 2006; Kamhi et al. 2010), we also found ini-tRNA among ALYREF interacting partners (**Table 3**, **Figure 6A** and Supplementary **Figure S3D**). Considering the importance of U5 snRNA in the spliceosome and the abundance of binding sites near the 5’SS, our findings align with ALYREF’s role in splicing and in the choice of LSSs.

**Figure 6.**
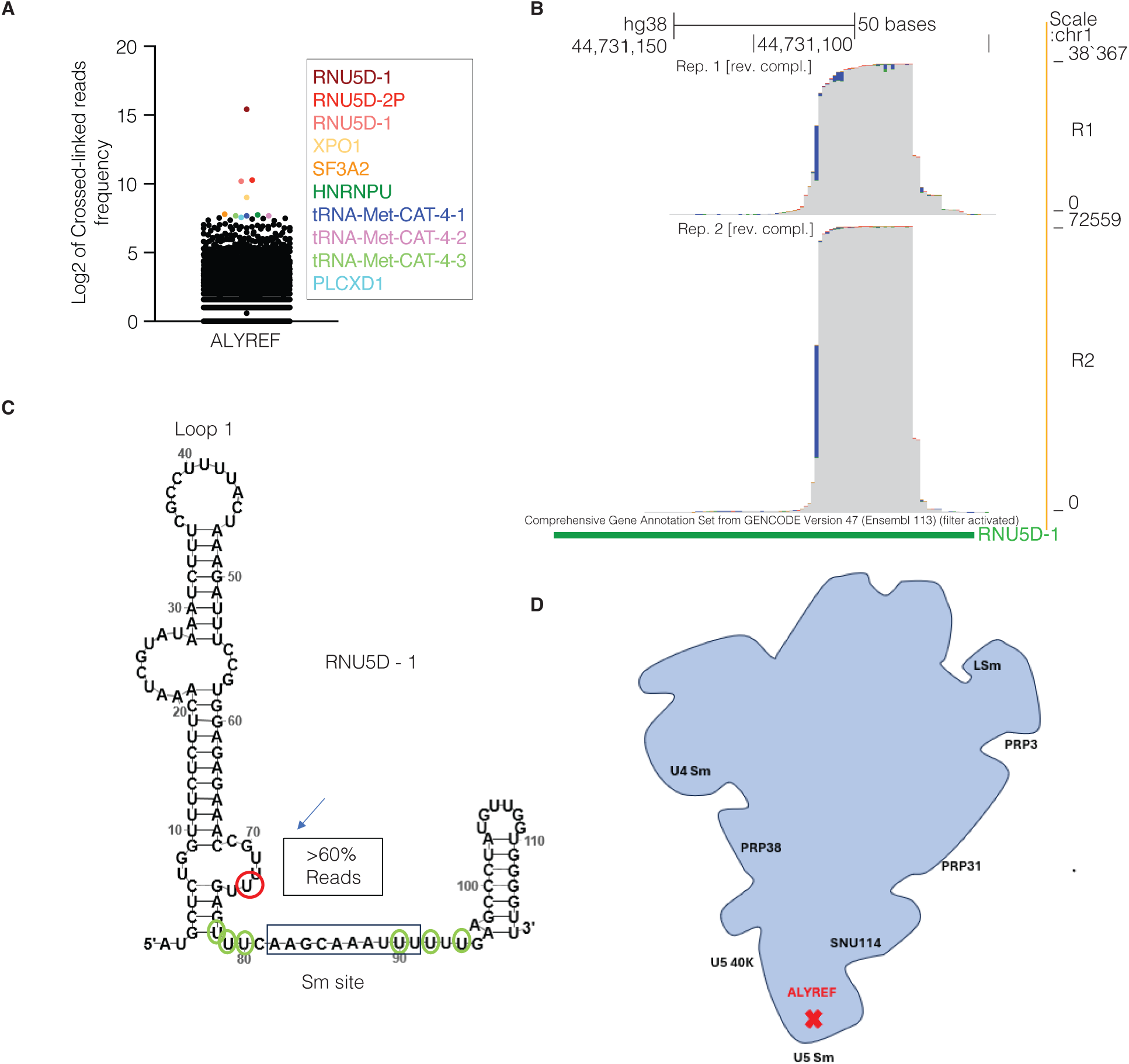
U5 snRNA is a top binding target of ALYREF. (**A**) RNU5 identified as top cross-linked RNA clusters in ALYREF fPAR-CLIP ranked using an in-house tool. (**B**) Alignment of nuclear endogenous ALYREF fPAR-CLIP reads on RNU5D-1 IGV tracks, illustrating binding sites. (**C**) Secondary structure of RNU5D-1, with top T-to-C changes (red circle, whereas green circles indicate additional changes). (**D**) A scheme of the spliceosome depicting the fPAR-CLIP proposed binding site of ALYREF.

**Table 3.** Summary of activated LSSs in samples with specific siRNA targets relative to two control samples. JI – Jaccard index (%).

| Control | AA5 | AA6 | AA5 & AA6 (JI) |
| --- | --- | --- | --- |
| Targeted factor |  |  |  |
| ALYREF | 833 | 785 | 466 (40.5) |
| PPIE | 1025 | 954 | 568 (40.3) |
| DDX41 | 1120 | 1121 | 625 (38.7) |
| DHX38 | 1081 | 1038 | 684 (47.7) |
| HNRNPA2B1 | 1252 | 1195 | 737 (43.1) |

### ALYREF deletion mutants abrogate SOS

To further investigate the role of ALYREF in SOS and LSS activation, we generated vectors that express HA-tagged full-length (WT) ALYREF as well as two deletion mutants: HA-ALYREF Δ130-150 and HA-ALYREF Δ195-240 (**Figure 7A**). First, we examined how these deletion mutants affect the activation of latent splicing in the wild-type CAD minigene construct (Li et al. 2002), which formed the basis of our luciferase reporter platform. Co-transfection of the CAD WT minigene with the ALYREF expression constructs showed that both deletion constructs activated latent splicing in the CAD minigene (**Figure 7B**). Next, we assessed the impact of both the WT ALYREF and the deletion mutants on the activation of latent splicing in the top activated LSSs after ALYREF knockdown, specifically BRD8, GPC5, ACADM, and LARS (**Figure 4**). Transfection with the HA-ALYREF Δ130-150 and HA-ALYREF Δ195-240 deletion mutants resulted in the activation of latent splicing in these four gene transcripts, compared to transfection with the WT ALYREF (**Figure 7C**). These overexpressed ALYREF mutants likely compete with the endogenous ALYREF for binding to the spliceosome. However, the deletions lack specific sequences required to function as SOS factors. These experiments confirmed that ALYREF is essential for SOS and suggest that amino acids 130-150 and 195-240 of ALYREF are important for SOS activation, potentially through binding to spliceosome components, altering subcellular localization patterns, or influencing the folding of ALYREF to create a functional protein involved in SOS (Supplementary **Figure S4**).

**Figure 7.**
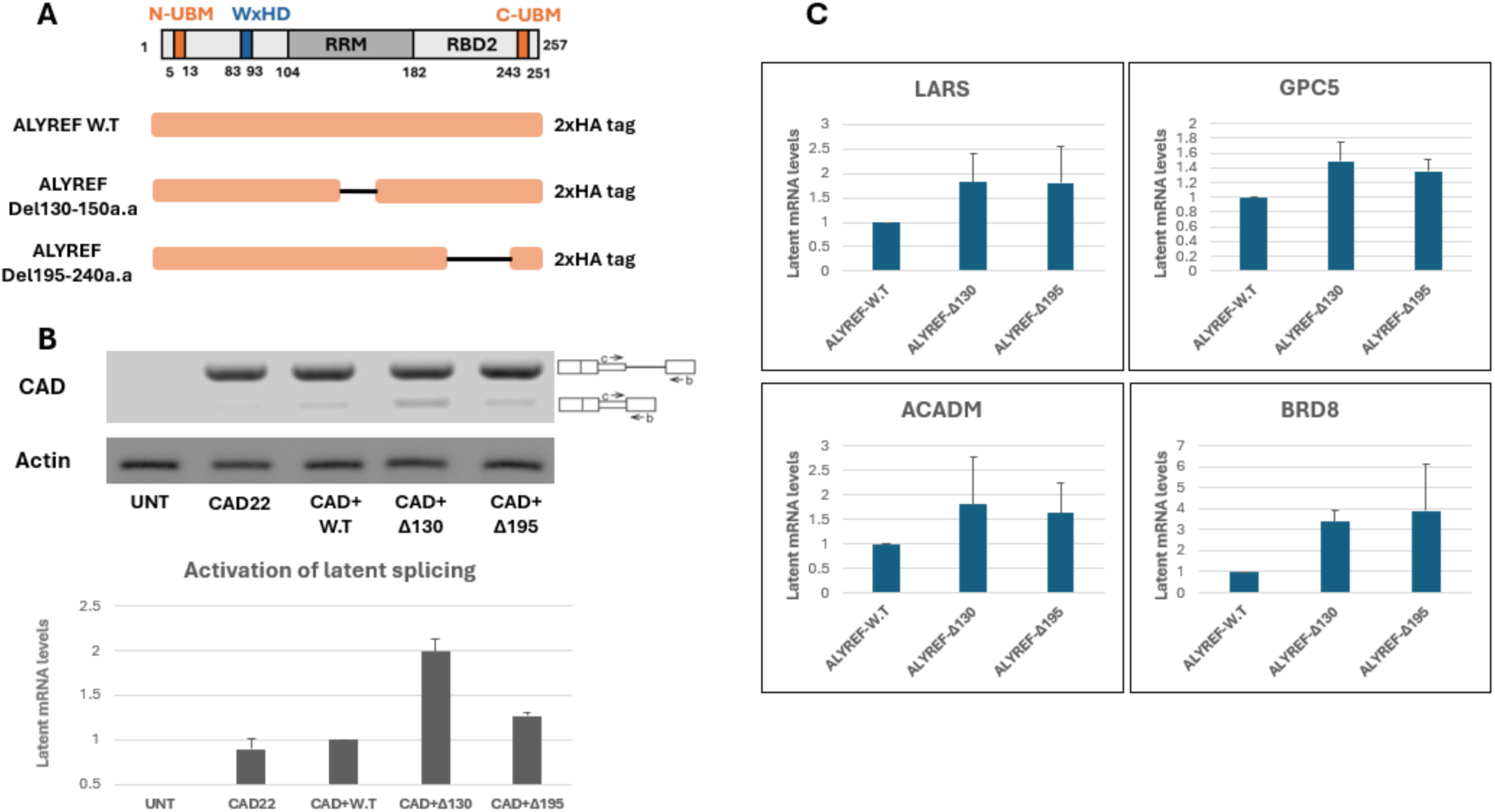
ALYREF mutants activate latent splicing. **(A)** Scheme of ALYREF domains and the WT and deletion mutants used. (**B**) RT-PCR analysis of latent splicing activation in the CAD WT minigene, when transfected into cells with constructs expressing either WT or each of the indicated ALYREF deletion mutants. (**C**) RT-PCR analysis of the effect of WT and each of the indicated ALYREF deletion mutants on activation of latent splicing of the top gene transcripts that show activation of latent splicing when ALYREF is knocked down (**Figure 4**).

### ALYREF mutants differ in their association with U5 snRNP and spliceosome components – Affinity purification studies

To confirm the association of ALYREF with U5 snRNA, we isolated nuclei from cells that were transfected for 24 hours with the HA-ALYREF construct. We then affinity-purified the nuclear complexes associated with transfected HA-ALYREF and confirmed the expected specific recovery of the transfected protein along with spliceosomal components, including the Sm protein (**Figure 8A**). Immunoblots with spliceosome components further showed co-immunoprecipitation of HA-ALYREF with spliceosomal proteins (Sm, PRP8, SNU114, SNRNP40) and the cap-binding protein CBP80 (**Figure 8A,B**). RT-PCR analysis confirmed the specific association of U5 snRNA with the transfected HA-ALYREF (**Figure 8C**). Similar experiments with ALYREF deletion mutants revealed that U5 snRNA also associates with these mutants, albeit to varying degrees (**Figure 8C**). Moreover, this analysis indicated that while the binding of the ALYREF deletion mutants to the Sm protein is only slightly reduced compared to the WT protein, binding to U5 snRNA-related proteins, namely, PRP8, SNU114, and SNRNP40, is negatively affected, especially by HA-ALYREF deletion Δ195-240 (**Figure 8B**). Similar results were also obtained after 48 hours of transfection (Supplementary **Figure S5**). Taken together, our functional assays (**Figure 7**) and other results indicate that ALYREF’s ability to suppress latent splice site activation depends on specific interactions with U5 snRNP components. Thus, ALYREF mediates SOS via the combined action of its RRM and RBD2 domains, which are required for engaging the U5 snRNA and its associated proteins.

**Figure 8.**
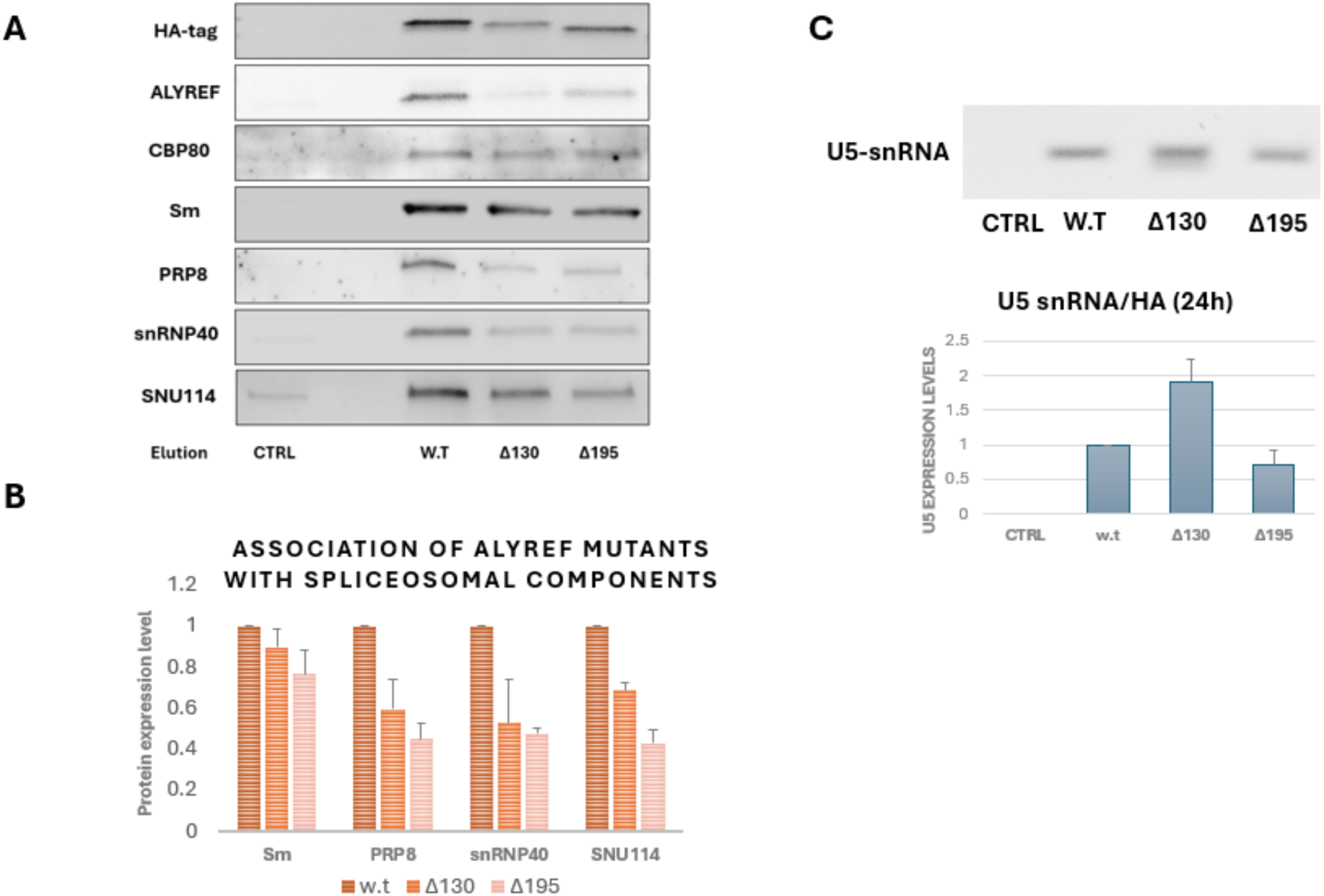
ALYREF mutants differ in their association with U5 snRNP and spliceosome components. **(A, B)** Analysis of affinity purified complexes assembled on nuclear WT HA-ALYREF and mutants (24-hour transfection). (**A**) WB. (**B**) Relative association of spliceosomal components with ALYREF mutants compared to the WT protein. (**C**) RT-PCR analysis of ALYREF-associated U5 snRNA (normalization of WT and control using Sm).

## Discussion

### Five RNA binding proteins as potential novel SOS factors

Splicing and AS regulation require the participation of a large number of RBPs (Kelemen et al. 2013; Akerman et al. 2015; Fredericks et al. 2015; Papasaikas and Valcarcel 2016; Fiszbein and Kornblihtt 2017; Sperling 2017; Dvinge 2018; Rogalska et al. 2024). This includes splicing proteins having an RNA recognition motif (RRM), which bind to the pre-mRNA and the U snRNAs. Splicing also involves a number of DExD/H helicases essential for the dynamic changes during the splicing reaction that remodel the spliceosome, as well as for proofreading (De Bortoli et al. 2021; Strittmatter et al. 2021). We, therefore, reasoned that RBPs are likely candidate SOS factors (Kamhi et al. 2010). Through a targeted siRNA screen directed against RBPs using a luminescence-based reporter platform, we identified ALYREF, PPIE, DHX38, and DDX41 as potential SOS factors. Future work with other reporters and screening conditions might identify more RBPs involved in SOS. Using the IRS vectors, we also identified hnRNPA2B1 as a potential SOS factor. In the screen, we could also confirm our previous observations that SOS is not dependent on NMD (Wachtel et al. 2004; Nevo et al. 2015) by specific knockdown of UPF1.

### An automated approach for detecting activated LSSs using RNA-seq

As described previously, identifying activated latent splice sites (LSSs) from RNA-seq requires both detection of noncanonical splice junctions and interpretation of surrounding expression patterns. While our previous work incorporated manual curation to remove artifacts and ambiguous events (Shefer et al. 2022), the scale of the current dataset necessitated a reproducible, automated pipeline. We therefore developed a filtering strategy based on relative expression across exonized regions, which yielded slightly more conservative results than manual inspection but remained consistent in overall significance and LSS profiles (Table S4). Importantly, this approach allows robust comparison across knockdown conditions and can be applied to future perturbation studies. Although LSS activation, as defined here, has not been systematically assessed in other large-scale splicing datasets, related phenomena such as aberrant exonization, intron retention, and noncanonical junction use have been reported in various contexts. For example, ENCODE Consortium RBP knockdown experiments (Van Nostrand et al. 2020) identified widespread splicing changes, though activation of latent sites per se was not explicitly characterized. Rogalska et al. (Rogalska et al. 2024) showed that knockdown of core spliceosomal proteins in HeLa cells induces substantial alternative splicing changes, including novel exon usage and intron retention. Their work highlighted the regulatory potential of the core spliceosome itself, revealing distinct AS patterns tied to specific complexes and subcomplexes. However, these studies did not explicitly characterize latent splice sites or examine their suppression as a conserved regulatory mechanism. Our findings build on this foundation by identifying specific RBPs that act as SOS factors to repress LSS activation and by dissecting the domain-level requirements for spliceosomal engagement. These data provide a mechanistic framework for understanding how splicing fidelity is maintained and how its disruption leads to the widespread activation of latent junctions. Cross-study comparisons are challenging due to differences in cell lines, sequencing depth, and annotation frameworks, but our observation of ∼5,000 reproducibly activated LSSs aligns with reported ranges of misspliced junctions in response to splicing factor depletion e.g., (Boutz et al. 2015; Dvinge 2018; Rogalska et al. 2024). Future studies could further refine these estimates by reanalyzing public RNA-seq data using our SOS-focused annotation framework.

Another consequence of this automated procedure is the identification of a higher number of activated LSSs compared with our previous study, even when considering only those selected in comparisons against both AA5 and AA6 controls (**Table 3**). This is likely also caused by the use of only one case replicate here. In contrast, we had two in our previous study, which reduced the stochastic variability associated with quantifying rare transcripts through RNA-seq experiments. While this aspect has less impact in determining whether factor knockdown results in increased LSS activation, since all four values used in Fisher’s exact test are affected, it has a greater impact on which specific LSSs are identified as activated. As a result, direct interpretation of which LSSs are activated should be approached with caution and should ideally be performed only after further curation of these datasets. Nonetheless, it is remarkable that the analysis of activated LSSs detected in more than one sample reveals distinct patterns that separate samples with specific siRNA targets from controls (**Figure 4B,C**), suggesting shared functional consequences of knocking down these five factors.

We also note here that the sequencing depth for our experiments (∼100 million reads) is only about half that of our previous study (Shefer et al. 2022). To investigate the impact of reduced sequencing depth on the detection of LSS activation, we compared our results with those obtained with even shallower datasets. Our comparison to RNA-seq data from DHX38 knockdown (Obuća et al. 2022) indicated that *i*) the depth of sequencing is important for detecting significant LSS activation, and *ii*) a sequencing depth of ∼100 million reads is sufficient for detecting LSS activation (see Suppl. Material File).

### ALYREF, PPIE, DHX38, DDX41 and hnRNPA2B1 identified as potential SOS factors

The sequencing results followed by the bioinformatics analysis revealed that hundreds of latent splice sites were activated upon knockdown of each of the five potential SOS factors (**Table 3**). These results were further validated for ALYREF, PPIE, DHX38, and HNRNPA2B1 by RT-PCR analysis of top transcripts that showed latent splicing activation (RNA-Seq results) after knockdown of the relevant potential SOS factor. These results portray ALYREF, PPIE, DHX38, and hnRNPA2B1 as novel SOS factors. We further demonstrate by RNA-seq that knockdown of each of these five splicing-associated factors results in increased usage of alternative donor splice sites that introduce in-frame STOP codons, thus suggesting they are functionally involved in controlling the selection of splice sites. These findings implicate ALYREF, PPIE, DHX38, and hnRNPA2B1 as functionally important in suppressing aberrant splice site usage. Although each has previously been linked to RNA processing, our results newly demonstrate that knockdown increases usage of 5′ latent splice sites that introduce premature stop codons. This suggests that diverse RNA-associated proteins, such as export factors, isomerases, helicases, and hnRNPs, may cooperate to preserve splicing fidelity beyond their canonical roles in spliceosome assembly or catalysis.

Recent molecular and structural studies identified the helicase DHX35 together with its cofactor, the G-rich protein GPATCH1, involved in a quality control mechanism to disassemble aberrant spliceosome intermediates. When catalytically active spliceosomes assemble with an aberrant 5’ splice-site conformation, they are rejected before the first catalytic step, by the helicase DHX35-mediated disruption of the U2 snRNP-branch site interaction, aided by GPATCH1. Meanwhile, the disassembly helicase DHX15, involved in spliceosome recycling, is found bound to the 3’ end of its U6 snRNA substrate (Li et al. 2025; Soni et al. 2025). In view of the important role of helicases in splicing, but also in quality control, it is interesting to note that among the five RBDs we identified as SOS factors, we identified two helicases.

### ALYREF a component of the supraspliceosome and a novel SOS factor

We chose to focus here on ALYREF because it was one of the two top hits of our screen for SOS factors and because it is a component of the supraspliceosome [(Kotzer-Nevo et al. 2014) and (Supplementary Figure S2)]. ALYREF is a conserved, multifunctional protein and a component of the TREX protein complex regulating the nuclear export of mRNAs (Masuda et al. 2005; Chi et al. 2013; Shi et al. 2017; Bonneau et al. 2023; Pacheco-Fiallos et al. 2023). It is recruited to intron-containing mRNAs by the splicing process (Masuda et al. 2005; Cheng et al. 2006; Taniguchi and Ohno 2008; Gromadzka et al. 2016; Viphakone et al. 2019; Pacheco-Fiallos et al. 2023), and by the cap-binding proteins (Cheng et al. 2006). Yet, it can also bind intron-less mRNAs (Taniguchi and Ohno 2008). Mutations or deletions in ALYREF disrupt interactions with the RNA and affect the interaction with the CBC (Cap-binding Complex) and EJC (Gromadzka et al. 2016). ALYREF can promote EJC multimerization, yet mutations in its WxHD and RRM domains impair the formation of the ALYREF-EJC-RNA and its multimerization *in vitro* and reduce binding to the mRNA (Pacheco-Fiallos et al. 2023). Mutations or deletions in ALYREF lead to changes in transcription and export defects, and ALYREF is further known to play a role in several types of cancer (Dominguez-Sanchez et al. 2011; Klec et al. 2022; Zhao et al. 2024). The main known role of ALYREF is in mRNA export from the nucleus to the cytoplasm through binding to the EJC (Shi et al. 2017; Viphakone et al. 2019; Bonneau et al. 2023; Pacheco-Fiallos et al. 2023). It should be noted that NMD plays no role in SOS, even though ALYREF is a core component of the EJC (which is crucial for recruiting UPF1 during NMD). Although indications for ALYREF involvement in splicing were reported (Viphakone et al. 2019), its role in latent splice site suppression is not fully understood.

Our fPAR-CLIP analysis of ALYREF revealed binding at canonical exon junctions, consistent with its established role in mRNA export (Viphakone et al. 2019). Notably, we also find binding across internal exon regions (**Figure 5E**) and at junctions of latent splice sites that become activated upon ALYREF knockdown (**Figure 5F**). These findings suggest that ALYREF directly engages with latent splice sites in vivo, supporting a model in which ALYREF acts co-transcriptionally to suppress aberrant splicing events.

### ALYREF binding to U5 snRNA - a major spliceosome component

Analyses of the binding partners and sites of ALYREF within the spliceosome, using fPAR-CLIP, revealed U5 snRNA as the top target exhibiting the most crosslinking signal at U73, which is located close to the Sm binding site of U5 snRNA. This finding establishes ALYREF as part of the active spliceosome (**Figure 6**). Affinity purification of nuclear complexes assembled on overexpressed ALYREF confirmed the association of WT ALYREF with U5 snRNA, together with splicing factors (**Figure 8**). However, ALYREF mutants lacking either 20 amino acids of the RBD (Δ130-150) or lacking part of the carboxy-terminus (Δ195-240) exhibit impaired function in SOS (**Figure 7**). These mutants also show impaired binding to U5 snRNP proteins (**Figure 8**). This finding indicates that the missing residues are relevant for the SOS function, either as being part of the binding sites to the U5 snRNA, to other spliceosome components, to the pre-mRNA, or through affecting the structure of ALYREF. Future experiments will be needed to dissect the specific role of ALYREF–U5 interactions in depth.

### ALYREF update of the SOS speculative model within the supraspliceosome

The endogenous spliceosome – the supraspliceosome is where splicing and AS, as well as all nuclear processing events of Pol II transcripts, occur in the cell nucleus. This huge (21 MDa) and highly dynamic splicing machine is composed of four active native spliceosomes, which are connected by the pre-mRNA (**Figure 9A**) (Sperling and Sperling 2017; Sperling 2017; Sperling 2019). The entire repertoire of nuclear pre-mRNAs, independent of their length and number of introns, is assembled in splicing-active supraspliceosomes, indicating their universal nature. The tetrameric structure of the supraspliceosomes is suitable to coordinate the multiple processing events of the pre-mRNA. Furthermore, it offers coordination and regulation of pre-mRNA processing events, including the SOS quality control mechanism (Sperling and Sperling 2017; Sperling 2017; Sperling 2019; Arafat and Sperling 2022).

**Figure 9.**
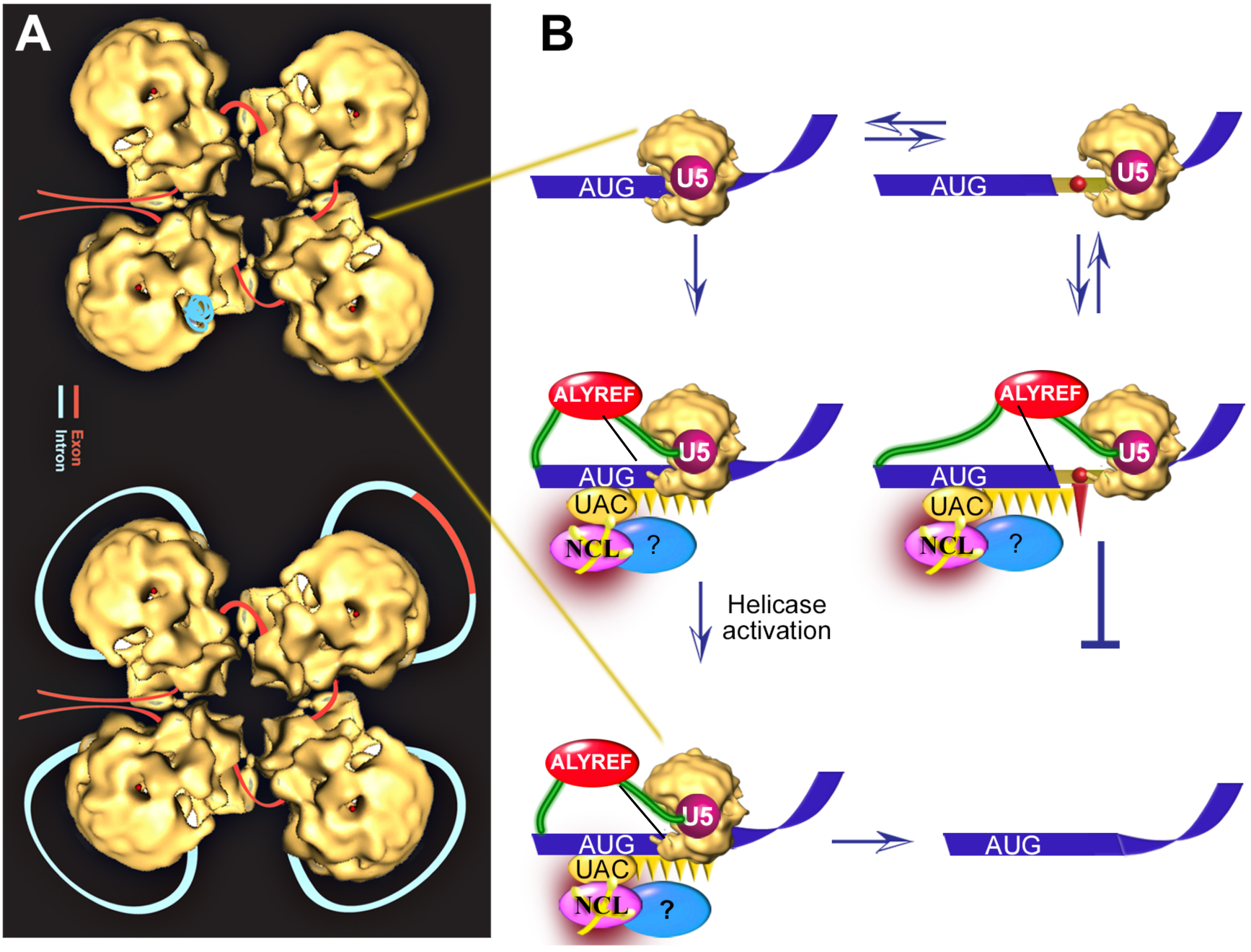
An updated speculative SOS model (Sperling 2017). (**A**) The supraspliceosome (Sperling and Sperling 2017; Sperling 2017; Sperling 2019). Exon, red; intron, light blue. (Top) The folded pre-mRNA that is not being processed is protected within the cavities of the native spliceosome. (Bottom) When a staining protocol that allows visualization of nucleic acids was used, RNA strands and loops were seen emanating from the looped-out scheme containing an alternative exon is depicted in the upper right corner. **(B**) Zoom into one spliceosome. Left scheme, splicing at the authentic 5**′**SS; right scheme, splicing at the latent 5**′**SS. Blue stripes, exons; grey line, intron; grey narrow stripe, latent exon; red circle, in-frame stop codon; gold, native spliceosome; orange ellipse (UAC), ini-tRNA; purple ellipse, NCL directly bound with ini-tRNA; and blue ellipse, additional associated components; orange triangles, hypothesized triplet-binding proteins; red triangle, stop-codon-binding protein; red ellipse, ALYREF binding to U5 snRNA (determined by fPAR-CLIP), as well as to CBC and EJC. Updated from refs.(Kamhi et al. 2010; Shefer et al. 2022).

The earlier proposed SOS working model (Kamhi et al. 2010) invokes a speculative sense triplet-recognition mechanism that can be interrupted by stop codon-binding proteins. We identified ini-tRNA as playing a pivotal role in SOS (Kamhi et al. 2010) and NCL, which is directly associated with ini-tRNA as the first identified protein component of SOS (Shefer et al. 2022).

Here, we established ALYREF as a novel SOS factor that binds to U5 snRNA and is required for SOS function. Notably, the fPAR-CLIP analysis here identified ini-tRNA, an essential component of SOS (Kamhi et al. 2010), as a binding target of ALYREF. With these results, we updated our working model of SOS as a quality control mechanism within the supraspliceosome that acts before splicing (**Figure 9**). Within the supraspliceosome, SOS approves the right combinations of splice junctions (**Figure 9B**). The first step of the SOS mechanism is the recognition of the AUG sequence by the complementary anticodon (UAC) of the initiator tRNA, which is in a complex with auxiliary proteins (Kamhi et al. 2010), including NCL, which is directly bound to the ini-RNA (Shefer et al. 2022). This step helps establish a register for the recognition of the reading frame. We now add ALYREF as an SOS factor through multiple binding targets within the supraspliceosome, including the pre-mRNA and mRNA, the CBC and EJC, but also through binding to U5 snRNA, a major spliceosome component placed at the heart of the spliceosome. We also identify a collection of RBPs newly implicated in SOS.

Further studies will be required to decipher the mechanism of SOS and determine the detailed role of the different factors involved. Namely, if SOS protein factors recognize specific sequence motifs on the pre-mRNA, or on the spliceosomal U snRNA, or some play a role through protein–protein interaction, or both, as the above speculative model predicts.

## Materials and Methods

### Plasmids

CAD WT minigene construct (Syrian hamster CAD) was previously described (Li et al. 2002). A luminescence-based reporter system IRS1 (Schuler et al. 2006) construct was prepared by cloning a luminescence-based reporter system based on a CAD minigene system that has been established and extensively studied in our group. It contains a latent 5’SS in its second intron, which has been repeatedly shown to be under SOS regulation. The reporter was constructed in the following manner: the first three exons and the two introns of the CAD minigene were used, and the first 40 nucleotides (1-40) of the cricket paralysis virus IRES (CrPVI) was inserted immediately upstream to the latent 5’SS in the second intron. The remaining sequence of CrPVI (nucleotides 41-190) was cloned immediately downstream of the 3’SS of the second intron, at the beginning of the third exon. This construct was cloned into the psiCHECK2 vector such that the Renilla luciferase gene would be expressed only when latent splicing has occurred to generate the full-length CrPVI IRES. This reporter construct was verified and its efficacy assessed ex-vivo by transfection experiments in which latent splicing was elicited by stress conditions and/or point mutations known to abrogate SOS. Two positive controls, IRS2 and IRS3, carry point mutations that change the translation start codon from ATG to ACG and AAA, respectively. Pre-mRNAs expressed from these mutants are expected to give a Renilla luciferase signal, because the mutations they carry have been previously shown to elicit latent splicing (Kamhi et al. 2006; Kamhi et al. 2010). The third mutant, IRS6, which was used as a negative control, carries a CAG to GCA mutation at the 3’ splice site used for both authentic and latent splicing, thus inactivating both splicing events. The four reporter constructs were verified by DNA sequencing, and quantitative estimates of latent splicing were made using luminescence as described above. Expression vectors of WT and deletion mutants of ALYREF WT, Δ130-150 and Δ195-240 (EX-H9330-M45; 130-150_EX-H9330-M45, and 195-240_EX-H9330-M45, respectively) cloned into the pReceiver-M45 Expression clone, having an HA tag, were purchased from GeneCopoeia.

### Screen of a library of siRNA against RNA binding proteins

First, the four luminescence IRS vectors were analyzed in a 96-well microplate. Forty-eight hours post-transfection into HEK 293T, using the Dual Glo Luciferase Assay System (Promega) (**Figure 1**). Next, the luminescence vectors were used to screen a library of siRNA pools for the knockdown of 766 RNA binding proteins (RBDs) in human cells. Each pool of siRNA consists of a mix of 4 different potent on-target siRNAs arrayed in 384-well microplates, with each well containing reagents that are directed to only one gene. U2OS cells were transiently transfected by either IRS1 or IRS6, 24hr post-transfection cells were plated on a 384-well microplate that was prepared with a battery of siRNA targeting various RBDs. The plates were thereafter incubated for 72 hours, and the luciferase output was monitored using the Dual Glo Luciferase Assay System (Promega) according to the manufacturer’s instructions. The screen was carried out in duplicate, and the average normalized luciferase reading from IRS1 and IRS6 was calculated. A final filtration step was carried out by dividing the IRS1 reading by the control IRS6. This step controls for any change in the Renilla luciferase output that is not derived from the eliciting of latent splicing, because the 3’ SS in the IRS6 was mutated (see above). Genes were considered positive hits only when the luciferase increase was seen in the two screen repeats and the increase was (at least) more than one standard deviation above the average read of the entire microplate (**Table 1**). These hits were subsequently verified and validated by deconvolution, where the siRNA pool was deconvoluted and each individual siRNA from the pool was tested. Hit verification requires at least two distinct siRNAs against one gene showing increase in the luciferase reading output, avoiding any potential off-target effects (**Table 1**). The IRS1, IRS6 vectors were also used for the analysis of the effect of downregulation of a number of hnRNPs, using a 96-well microplate (**Figure 2**).

### Knockdown of selected gene transcripts

HEK293 or HeLa cells were individually transfected with siRNA (the relevant ON-TARGETplus Human siRNA SMARTpool, Dharmacon) targeted to each of the following gene transcripts: hnRNPA2B1 (L-011690-01-0005), ALYREF (L-012078-00-0005), PPIE (L-009466-01), DHX38 (L-013428-00-0005) and a non-targeting siRNA as control (D-001810-01-05) (Dharmacon). Cells were grown in 6-well plates and transfected with 45 nM siRNA using TransIT-LT1 Transfection reagent (Mirus) according to the manufacturer’s protocol. After 24 h, the medium was changed, and cells were transfected again, as previously described. After 48 h, total proteins and RNA were extracted (RNeasy Qiagen 74104) according to the manufacturer’s instructions and analyzed.

### Western blot (WB)

Samples were separated on 12% SDS–PAGE and transferred to a nitrocellulose membrane. The membrane was blocked with 5% BSA for 30 min at room temperature, following 1hr incubation at 4°C with the relevant primary antibody: hnRNPA2B1-ab31645; THOC4-ab202894; DHX38-ab154801 (Abcam, Tel Aviv, Israel), PPIE-sc-100700 (Santa Cruz Biotechnology), HA (C29F4, Cell Signaling) and GAPDH (G8796, Sigma, 1:1000). The secondary antibody was horseradish peroxide-conjugated goat anti-rabbit IgG and goat anti mouse (1:5000 dilution; H + L; Jackson ImmunoResearch) incubated for 15 min at room temperature. The membrane was visualized using SuperSignalTM West Femto Stable Peroxide (34095, Thermo Scientific) on the Fusion instrument (A2S).

### RNA analysis

RNA was extracted from cells, using RNeasy Qiagen kit (74104), and used to prepare cDNA. PCR was performed on the cDNA using the relevant primers (Supplementary Table S3). The PCR products were separated on 2% of agarose gel. The identity of all PCR products was confirmed by Sanger sequencing. Each experiment was repeated at least 3 times.

### Knockdown and global analysis by RNA Seq

HeLa cells were forward transfected in 96-well plates with an siRNA library (genome-wide screen, SMARTpools siGENOME, Thermo-Scientific) using an automatized robotic procedure (Sciclone Liquid Handling Workstation, Perkin Elmer). Cellular mRNAs were isolated 72 hr. post-transfection by an automated procedure using oligo-dT-coated 96-well plates (mRNA catcher PLUS, Life Technologies) following the manufacturer’s recommendations. Control samples included five non-targeting siRNA and two mock transfected samples. The siRNA knockdowns were sequenced with the following parameters: 100 M reads pair ended, 126 bp length (Rogalska et al. 2024).

### RNA-seq data evaluation

RNA-seq reads were aligned with STAR against the human genome (hg19 assembly) as previously described (Shefer et al. 2022). A summary of aligned reads is provided in **Table 2** for five case samples, labeled by the name of the factor targeted by siRNA, five control samples (non-specific siRNA) labeled AA3 through AA7, and two negative controls (mock transfections), labeled AA8 and AA9. Evaluation of data quality and quantification of splice junction support with split reads was done with the QoRTs package (Hartley and Mullikin 2015) (Hartley and Mullikin). Three samples, AA3, AA7, AA8, were found to be affected by reduced diversity of mapped reads (Fig. S6A), and two of them, AA3 and AA8, were also found to exhibit increased 3’ bias of mapped reads (Fig. S6B). Therefore, these three samples were not included in further analyses.

### Identification of increased usage for junctions of interest

Evaluation of increased usage of junctions of interest (LSS, adSS3n) was done in a pairwise manner, with one sample considered the “case” and the other as the “control”. Using quantifications of split reads (i.e. junction reads) obtained with the QoRTs package, we computed junction usage relative to the canonical junction which contains the intronic sequence in which the junction of interest is located. We selected those junctions that exhibited an increase in relative usage of at least 50% between the case and control samples. Furthermore, for the junctions that passed this initial thresholding, we constructed expression profiles for the intronic regions exonized through the use of LSS or adSS3n junctions (i.e. exon extension) and evaluated them visually or automatically for consistency with expectations for exonized regions. Of note, in building expression profiles we eliminated reads mapping in the intronic region downstream from the LSS or adSS3n, in effect minimizing the impact of intronic transcriptional noise. The automated evaluation required that the “case” sample exhibited higher expression levels across 90% of the exon extension and 1.5-fold or higher expression level across 60% of the exon extension. This automated approach has the benefit that it eliminates human inconsistencies, but this is at the expense of not eliminating all false positives.

### Evaluation of SOS status

As described previously (Shefer et al. 2022), disturbance of the proper function of the SOS mechanism in specific samples can be inferred from an increase in the number of LSSs with elevated usage (i.e. activated). Here we applied the same method to evaluate the number of activated LSSs upon knockdown of specific factors. Specifically, we conducted a one-sided Fisher’s exact test (FET) with four quantities: LSSs activated in the “case” relative to “control” (i.e. activated), LSSs activated in “control” relative to “case” (i.e. de-activated), activated adSS_3n_, and de-activated adSS_3n_ (adSS_3n_ serve as proper controls since they do not lead to the inclusion of in-frame STOP codons). Visual evaluation of all expression profiles becomes prohibitive for this analysis due to the high number of junctions that pass the initial thresholding based on split read support of splice junctions, so we evaluated whether replacement of the visual with the automatic evaluation of profiles yields similar results. For this purpose, we conducted four comparisons (DDX41, DHX38, PPIE, THOC4 vs. AA4) where expression profiles were evaluated both visually and automatically as described above. In all four cases, the p-values associated with corresponding FETs were very similar (Table S4), with the “automated” evaluation yielding only slightly higher values (i.e. more conservative). This is likely due to the inability of the automatic evaluation to eliminate as many false positive cases as in the case where profiles are evaluated visually.

### Evaluation of activated LSS commonalities

To evaluate the significance of observed LSS activated upon knockdown of two separate factors, we conducted Monte Carlo simulations (ten thousand replicates) that allowed us to estimate the expected number of LSS activated in two separate samples. For this purpose, in each replicate, we randomly selected the same number of LSSs as found activated in each sample (**Table 3**) from the initial set of LSSs supported by at least four split reads (**Table 2**) and determined the number of common occurrences between samples in a pairwise manner. The expected values are reported as the mean of the distribution obtained for each pairwise comparison (**Figure 3**).

### Fluorescent Photoactivatable Ribonucleoside-Enhanced Crosslinking and **Immunoprecipitation (fPAR-CLIP).**

fPAR-CLIP for endogenous ALYREF was conducted based on the protocol outlined previously (Anastasakis et al. 2021), with specific modifications. HEK293 cells were seeded at 1.3×10^8^, and upon reaching 70% confluence, they were treated with 100 μM 4-thiouridine for 16 hours before being crosslinked at 0.4 J/cm^2^ under 365 nm UV light. Approximately 560 μL of nuclear supernatants enriched in supraspliceosomes were isolated as previously reported (Spann et al. 1989; Azubel et al. 2006). Immunoprecipitation was conducted using 20 μL anti-ALYREF antibody (Abcam; ab20894) coupled to 75 μL protein – G magnetic beads (Sigma). The beads were treated with 0.15 U/μL RNaseI, dephosphorylated, and ligated to a 3′-end fluorescent adapter labelled with Alexa Fluor 647 (5’-rAppNNTGACTGTGGAATTCTCGGGT(fl)GCCAAGG-*2xAF647*). Samples were separated using NuPAGE^TM^ 4-12% Bis-Tris SDS-PAGE (Invitrogen), and the gel was scanned at 647 nm using Typhoon FLA950 scanner. Fluorescent ALYREF-adapter bands were excised, shredded, and digested with proteinase K. RNA was isolated using phenol-chloroform extraction, and a cDNA library was generated using an RT primer (GCCTTGGCACCCGAGAATTCCA). A Low cycle PCR was performed using Platinum Taq DNA polymerase (Thermo Fischer Scientific, 10966034) with a short 5′ PCR primer (CTTCAGAGTTCTACAGTCCGACGA) and the RT primer according to the manufacturer’s instruction. PCR products were size-selected (75-100 bp) using a 3% agarose gel (Pippin Prep cassettes, Sage Science, CSD3010), followed by a large-scale PCR using an optimized 16-cycle protocol. The final PCR products were purified using the DNA Clean & Concentrator^TM^-5 kit (Zymo Research, D4013) and size-selected (∼135-175 bp) on a 3% Pippin Prep to eliminate primers and adapter-adapter byproducts. The TapeStation (Agilent) system was used to assess the library quality, and the resulting cDNA library was sequenced on an Illumina NovaSeq 6000 or NovaSeq XPLUS platform.

fPAR-CLIP data were processed using the PARpipe pipeline, incorporating PARAlyzer for peak calling. Within the pipeline, adapters were trimmed, and Bowtie1 was used to align FASTQ files to the human reference genome (hg38) with the following parameters: `-v 2 -m 2 --all --best --strata`. The resulting `clusters.csv` files provided comprehensive metrics for ranking and filtering clusters. To evaluate the reproducibility of Endo-ALYREF fPAR-CLIP replicates, deepTools was used to quantify aligned reads across uniform genomic bins. Normalized read counts were employed to compute Spearman’s correlation coefficient (ρ), enabling a robust comparison of replicate similarity.

To obtain profiles of the ALYREF binding, we overlapped the set of transcriptome regions of interest centered on specific features (e.g. 5’SS, LSSs) with the genomic coordinates of the read clusters as determined above. At each relative position within the region of interest we determined the overlap with a read cluster as a binary event regardless of how many reads were found in each read cluster. The final profile was constructed by plotting average occurrences of read clusters at each relative position across all regions of interest.

### Overexpression of ALYREF WT and deletion mutants

HeLa cells were grown in 6 well plates and transfected with 1 µg of each of the ALYREF expression vectors: WT, Δ130-150 and Δ195-240, using TransIT-LT1 Transfection reagent (Mirus) according to the manufacturers’ protocol. The three ALYREF expression constructs have two HA repeats in their carboxy terminus. 48 hr. post-transfection cells are collected using RLN buffer to separate nuclear and cytoplasmic fraction (Agranat-Tamir et al. 2014). The nuclear phase was diluted with ST2Mx1 buffer with protease inhibitor, and the samples were sonicated. For protein analyses 100 µl were used, while 900 µl were taken for RNA extraction using RNeasy (Qiagen 74104) according to the manufacturer’s instructions. cDNA was synthesized using dT15 primer, followed by RT-PCR.

For the co-transfection of ALYREF mutants with CAD minigene, HEK293 cells were grown in 6 well plates and transfected with 1 µg of CAD WT minigene, and 1 µg of each of the ALYREF expression vectors using PEI. RNA was extracted 24 hr. post-transfection, cDNA was synthesized using dT15 primer, and analyzed by RT-PCR

### Affinity purification of nuclear complexes associated with the HA-tagged ALYREF

HeLa cells were grown in 6-well plates, transfected with 1 µg of each of the different HA-tagged ALYREF expression vectors: WT, or one of the following deletions: Δ130-150 and Δ195-240, using the TransIT-LT1 transfection reagent (Mirus), according to the (Agranat-Tamir et al. 2014) manufacturer’s protocol. Cells were collected 48 or 24 hr. post-transfection using RLN buffer to isolate nuclear and cytoplasmic fractions (Agranat-Tamir et al. 2014). The nuclear fraction was aspirated with ST2M buffer and protease inhibitor, and the samples were sonicated. For affinity purification of nuclear complexes associated with the different ALYREF mutants, the nuclear samples were immunoprecipitated using HA magnetic beads (Pierce, 88836) for 30 min at 4°C with rotation. The beads were washed three times with ST2M buffer, followed by elution of the RNA and proteins from the beads. Proteins were eluted with 2XSDS sample buffer at 95 ^°^C for 5 min, and RNA was eluted from the beads as described (Azubel et al. 2006). Briefly, the beads or the nuclear samples were mixed with extraction buffer (50 mM Tris, pH 7.5, 150 mM NaCl) and 10% SDS. The RNA was recovered by extraction with phenol and precipitation in ethanol. Next, the RNA was treated with DNase I (50 U/ml; Promega), and cDNA was synthesized as described using dT15 primer (Shefer et al. 2022). WB with anti-Sm was used for normalization of untransfected and ALYREF WT-transfected cells. For analyses by RT-PCR of U5 snRNA associated with each of the transfected HA-ALYREFs, we used the level of U5 snRNA in untreated and ALYREF WT-transfected cells for normalization, which was further normalized to transfection efficiency as reflected by the level of the associated HA-tagged ALYREF.

## Supplementary Materials

**Supplementary Table S1.** RBP screen results

**Supplementary Table S2.** fPAR-CLIP results

**Supplementary Table S3.**
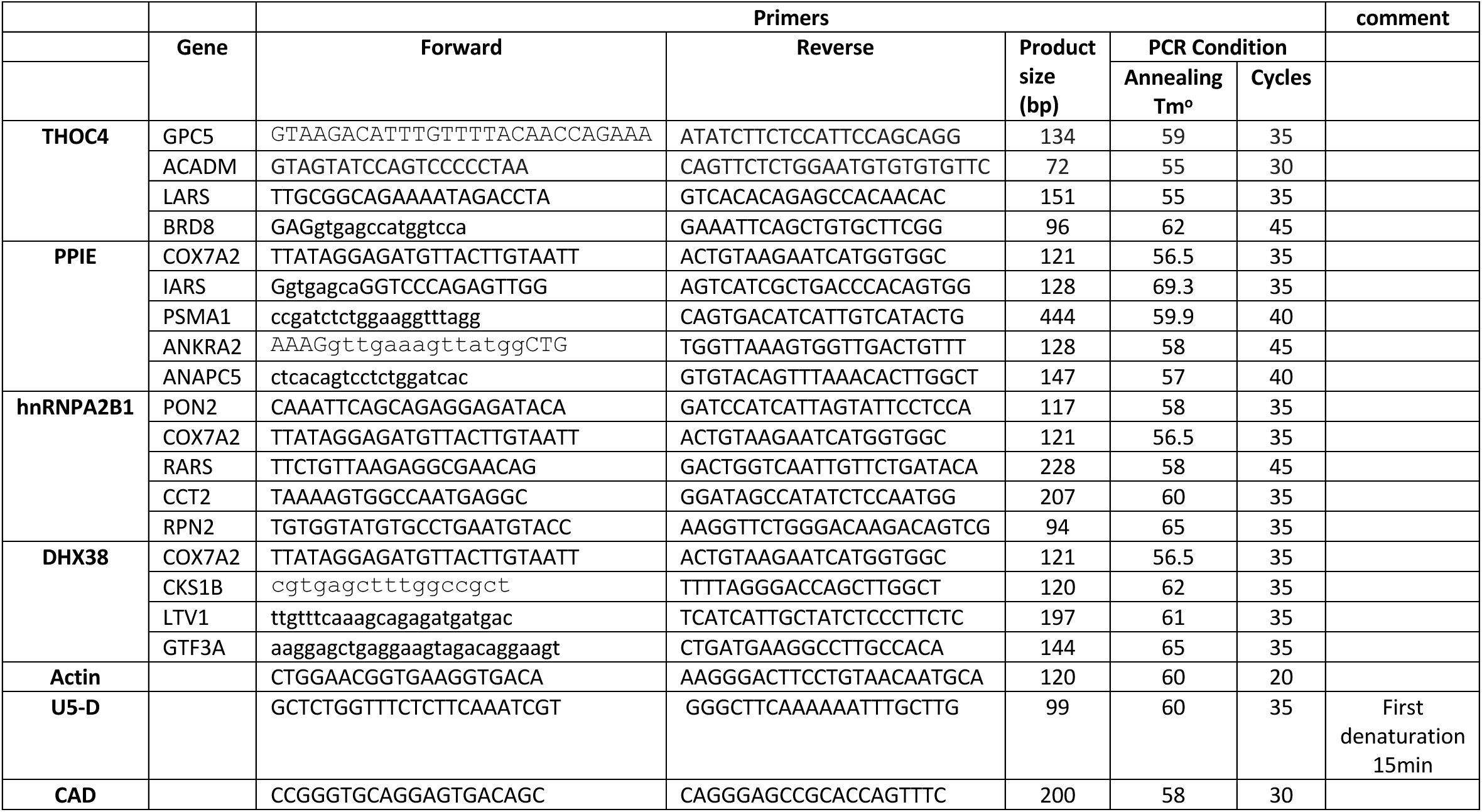
List of PCR primers

**Table S4.** Comparison of visual (V) and automated (A) evaluations of increased activation of LSSs upon specific factor knockdown (KD) relative to the AA4 control sample. adSS – alternative donor splice site; A.LSS – LSSs activated upon factor KD; D.LSS – LSSs de-activated (or activated in the control relative to the KD samples); A.adSS_3n_ – adSS_3n_ activated upon factor KD; D.adSS_3n_ – de-activated adSS3n; OR – odds ratio; p-val – p-value associated with one-sided Fisher’s exact test.

| adSS Type |  | A.LSS | D.LSS | A.adSS<sub>3n</sub> | D.adSS<sub>3n</sub> | OR | p-val |
| --- | --- | --- | --- | --- | --- | --- | --- |
| Targeted factor |  |  |  |  |  |  |  |
| ALYREF | V | 578 | 703 | 161 | 221 | 1.129 | 0.1665 |
|  | A | 803 | 1127 | 175 | 245 | 0.998 | 0.5316 |
| PPIE | V | 721 | 653 | 206 | 180 | 0.965 | 0.6435 |
|  | A | 934 | 932 | 227 | 190 | 0.839 | 0.9529 |
| DDX41 | V | 723 | 673 | 212 | 204 | 1.034 | 0.4046 |
|  | A | 1139 | 1069 | 245 | 221 | 0.961 | 0.6698 |
| DHX38 | V | 716 | 622 | 164 | 201 | 1.411 | 0.0022 |
|  | A | 1011 | 851 | 184 | 213 | 1.375 | 0.0023 |

## Funding

This study was partially supported by grants from the Israel Science Foundation [(ISF) 624/13; 1902/22], and the Israel Cancer Research Fund (ICRF) to R.S.

The Helen and Milton Kimmelman Center for Biomolecular Structure and Assembly at the Weizmann Institute of Science to J.S.

The European Research Council (Horizon ERC-2023-ADG-101141245) to R.A.

The Intramural Research Program of the National Institutes of Health, National Institute of Arthritis and Musculoskeletal and Skin Diseases (ZIA-AR041205 to M.H.)

The Intramural Research Program of the National Institutes of Health, National Human Genome Research Institute (ZIA-HG200323 to L.E.)

## Supporting information

SOS_Supplementary_FIGs and Materials

## Acknowledgments

We thank Aviva Petcho for excellent technical assistance. We thank Dr. Juan Valcarcel (ICREA and Center for Genomic Regulation (CRG), Barcelona, Spain) and his lab for making RNA-seq datasets available to us before publication. The authors would also like to thank Dr. Stefania Dell’Orso, Faiza Naz, and Shamima Islam (Genomic Technology Section, NIAMS) for support with high-throughput sequencing.

