## Supplementary material for "Identification of RNA binding proteins that mediate a quality control mechanism of splicing": SOS_Supplementary_FIGs and Materials

**Figure S1. Luciferase expression assay of latent splicing induced by heat shock.** HEK 293T cells were transfected with IRS1. Twenty-four hr post-transfection, the cells were heat-shocked at 45ºC for 45 min, returned to 37ºC and incubated for the indicated time length. Levels of latent splicing are expressed as the ratio between the Renilla luminescence to that of the Firefly luciferase, as measured by the Promega dual-luciferase system.

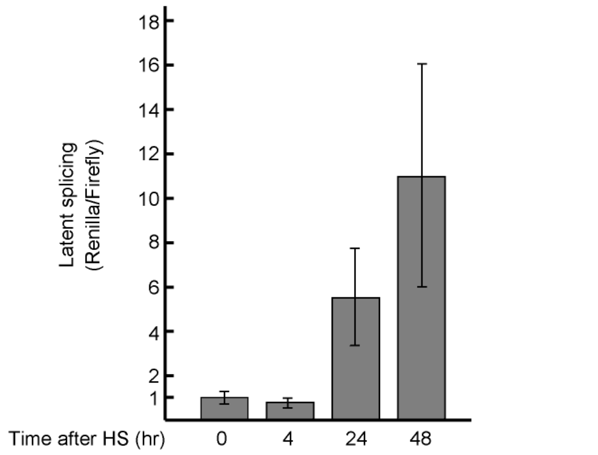


**Figure S2. Alyref is a component of the supraspliceosome.** (**A**) Schematics of isolation of supraspliceosomes from cells expressing PP7-tagged SMN1 minigene transcript as described(Spann et al. 1989; Azubel et al. 2006), followed by affinity purification of supraspliceosomes assembled on PP7-tagged SMN1 minigene transcript as described(Kotzer-Nevo et al. 2014). (**B**) WB of nuclear supernatants from cells expressing PP7-tagged SMN1 and control cells (left), and affinity purification of supraspliceosomes from the PP7-SMN1 expressing sample and control, revealing the association of ALYREF with supraspliceosomes.

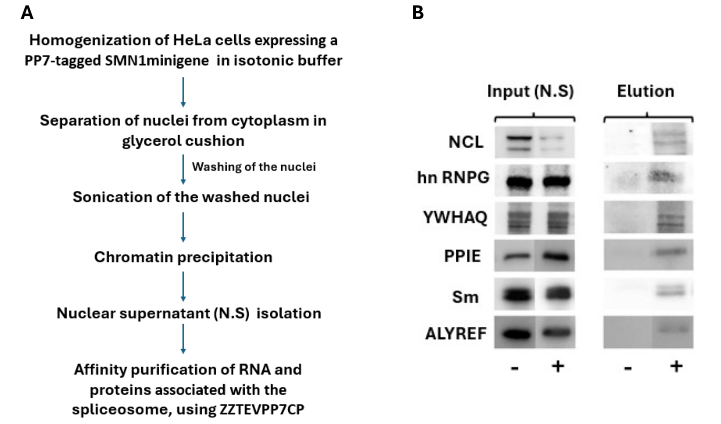

A

| Replicate | 1 | 2 |
|---|---|---|
| Total Number of clusters | 14,968 | 5,235 |
| No Annotation clusters | 794 | 202 |
| Annotated clusters | 14,174 | 5,033 |

B

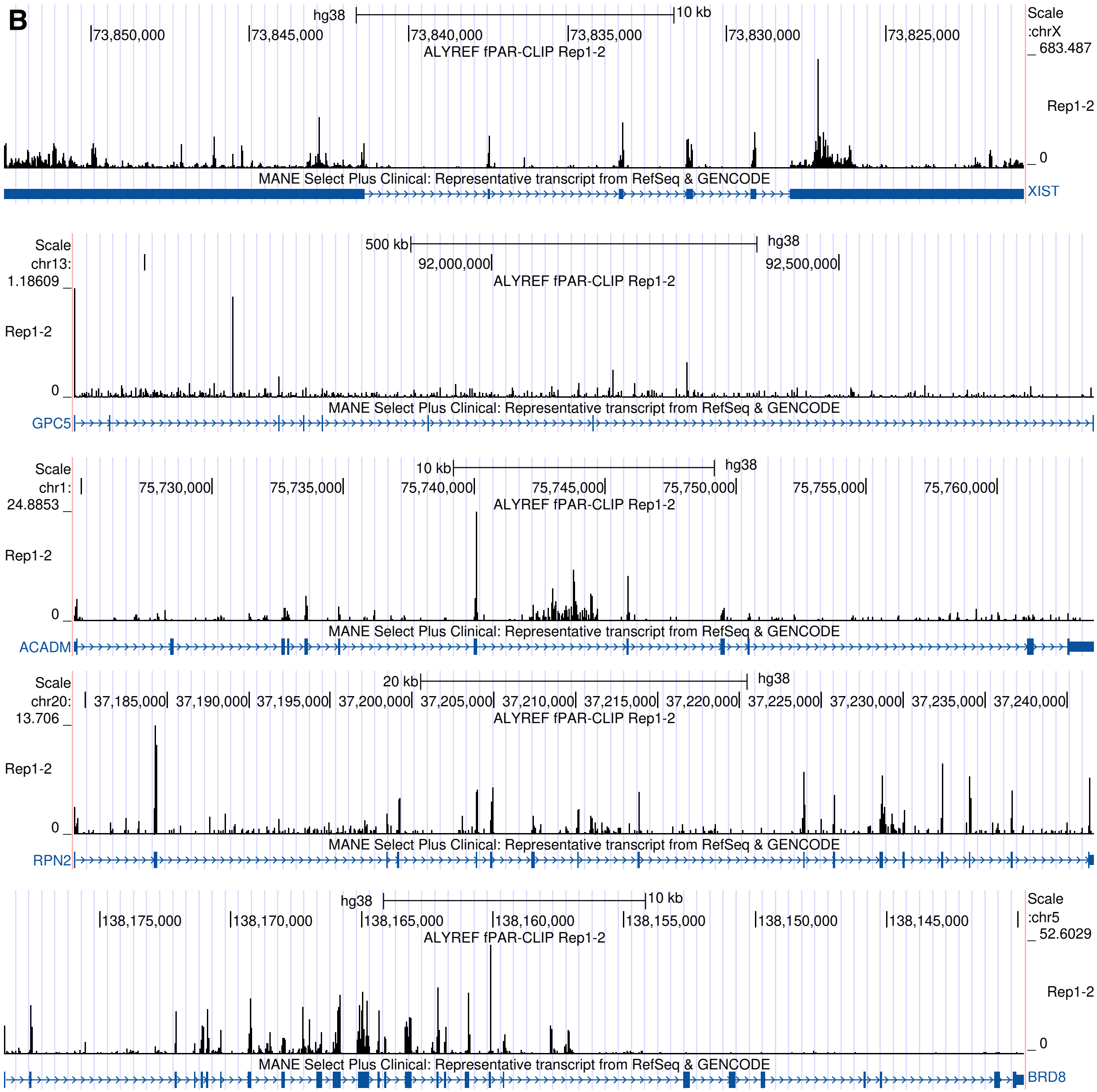

Scale
hg38
10 kb
chrX:
73,850,000
73,845,000
73,840,000
73,835,000
73,830,000
73,825,000
ALYREF fPAR-CLIP Rep1-2
683.487
Rep1-2
0
MANE Select Plus Clinical: Representative transcript from RefSeq & GENCODE
XIST
Scale
chr13:
500 kb
hg38
92,000,000
92,500,000
ALYREF fPAR-CLIP Rep1-2
1.18609
Rep1-2
0
MANE Select Plus Clinical: Representative transcript from RefSeq & GENCODE
GPC5
Scale
chr1:
10 kb
hg38
75,730,000
75,735,000
75,740,000
75,745,000
75,750,000
75,755,000
75,760,000
ALYREF fPAR-CLIP Rep1-2
24.8853
Rep1-2
0
MANE Select Plus Clinical: Representative transcript from RefSeq & GENCODE
ACADM
Scale
chr20:
20 kb
hg38
37,185,000
37,190,000
37,195,000
37,200,000
37,205,000
37,210,000
37,215,000
37,220,000
37,225,000
37,230,000
37,235,000
37,240,000
ALYREF fPAR-CLIP Rep1-2
13.706
Rep1-2
0
MANE Select Plus Clinical: Representative transcript from RefSeq & GENCODE
RPN2
Scale
hg38
10 kb
chr5:
138,175,000
138,170,000
138,165,000
138,160,000
138,155,000
138,150,000
138,145,000
ALYREF fPAR-CLIP Rep1-2
52.6029
Rep1-2
0
MANE Select Plus Clinical: Representative transcript from RefSeq & GENCODE
BRD8

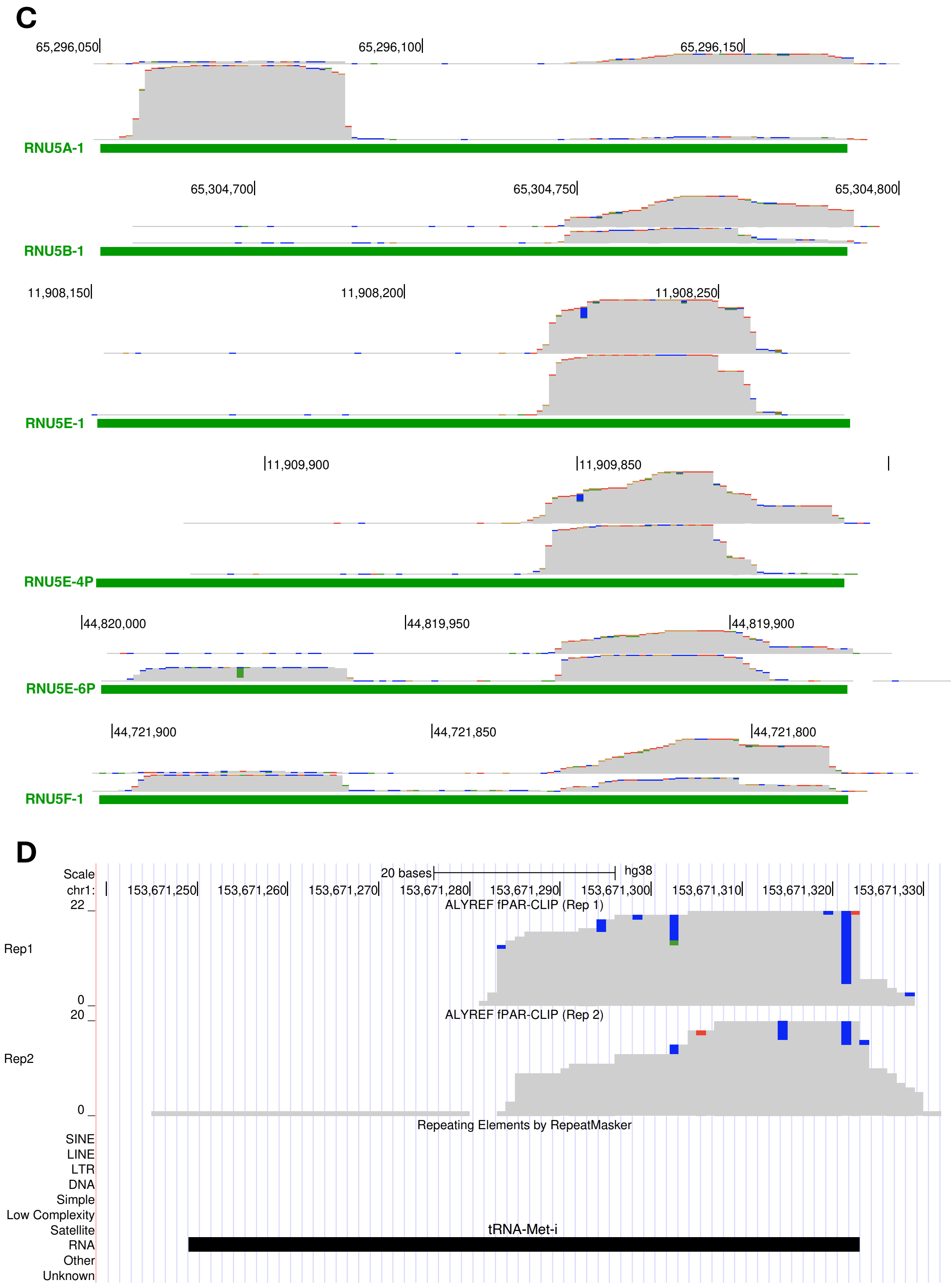
C
65,296,050
65,296,100
65,296,150
RNU5A-1
65,304,700
65,304,750
65,304,800
RNU5B-1
11,908,150
11,908,200
11,908,250
RNU5E-1
11,909,900
11,909,850
RNU5E-4P
44,820,000
44,819,950
44,819,900
RNU5E-6P
44,721,900
44,721,850
44,721,800
RNU5F-1
D
Scale
20 bases
hg38
chr1:
153,671,250
153,671,260
153,671,270
153,671,280
153,671,290
153,671,300
153,671,310
153,671,320
153,671,330
22
ALYREF fPAR-CLIP (Rep 1)
Rep1
0
20
ALYREF fPAR-CLIP (Rep 2)
Rep2
0
Repeating Elements by RepeatMasker
SINE
LINE
LTR
DNA
Simple
Low Complexity
Satellite
RNA
Other
Unknown
tRNA-Met-i

**Figure S3. Classification and Genomic Distribution of ALYREF fPAR-CLIP Clusters (A)** Classification of ALYREF fPAR-CLIP clusters into annotated and non-annotated regions. (**B**) ALYREF fPAR-CLIP pile-ups shown as density graphs across five gene loci, illustrating binding sites. Reads were combined from two replicate experiments. Loci are shown in the 5'-3' orientation of the gene. Gene names are shown next to the gene model. (**C**) Density graphs of read pileups at five U5 snRNA loci for two fPAR-CLIP replicates for each locus. Track heights are proportional to the height of the read pile-up, ranging from 1590 (RNU5A-1, replicate 1) to 11442 (RNU5A-1, replicate 2) reads. Green bars correspond to the U5 RNA as annotated in the GENCODE v47 collection. Loci are shown in the 5'-3' orientation of the U5 RNAs. Graph colors indicate read matches to hg38 consensus (gray), or mutations (A – green, C – blue, G – orange, T – red). (**D**) Density graphs of read pileups at one ini-tRNA locus on chr1 for two fPAR-CLIP replicate experiments. The location of the ini-tRNA is provided by the RepeatMasker annotation, as indicated by the black bar. Read colors are the same as in (**C**).

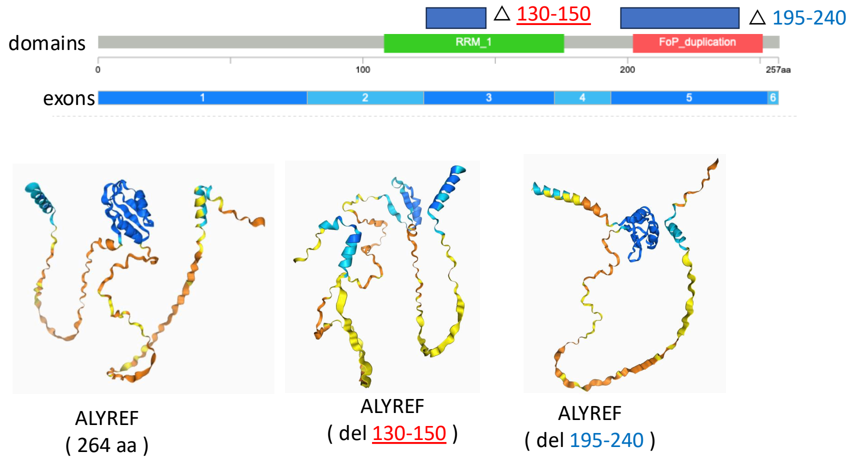


**Figure S4. Motives and predicted conformations of WT ALYREF and deletion mutants.** Structures were produced with AlphaFold 3.

**Figure S5. ALYREF mutants differ in their association with U5 snRNP (A, B).** Analysis of affinity purified complexes assembled on nuclear WT HA-ALYREF and mutants (48 hr transfection). (**A**) WB. (**B**) RT-PCR analysis of ALYREF-associated U5 snRNA (normalization of WT and control using Sm).

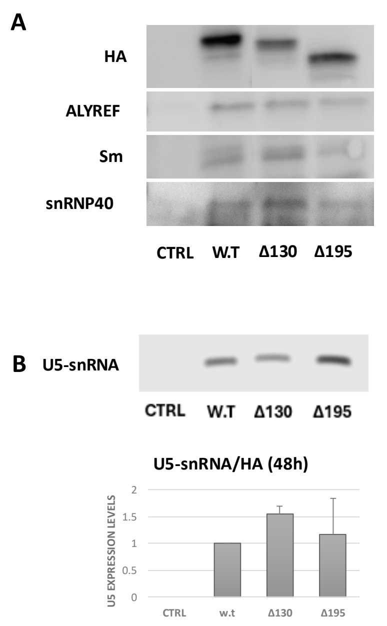


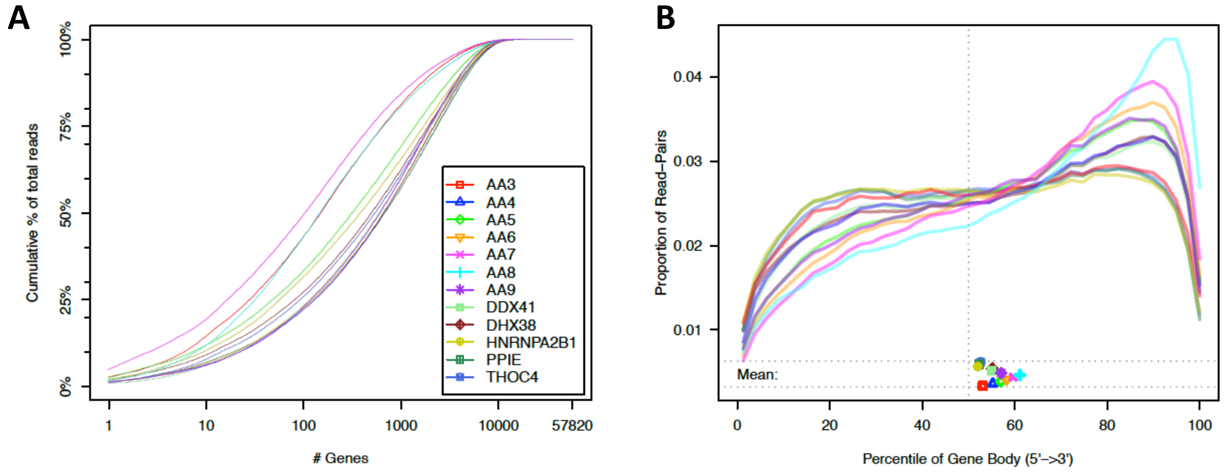


**Figure S6**. Select quality control measures of RNA-seq experiments provided by the QoRTs tool. (**A**) Cumulative gene assignment diversity of reads. Three samples (AA3, AA7, AA8) present lower diversity read assignments compared to the rest of the samples. (B) Gene body coverage calculated for genes within the upper middle quartile expression level range. Two samples (AA7, AA8) are the most affected by read mapping biased toward the 3' end of genes.

__Supplementary Material__

### Identification of RNA binding proteins that mediate a quality control mechanism of splicing

**Maram Arafat[1#], Valer Gotea[2#], Mubarak I Umar[3#], Aya Muharram[1], Eyal Kamhi[4,] Reuven Agami[5], Joseph Sperling[4*], Markus Hafner[3*], Laura Elnitski[2*], and Ruth Sperling[1*]**

**Knockdown of DHX38 in an independent experiment.**

Increase in public availability of data from other siRNA experiments allowed us to attempt a verification of the observations made herein with an independent dataset. In the context of retinitis pigmentosa, DHX38 was down regulated via RNAi with the purpose of identifying its impact on general splicing, as well as on splicing of retina specific genes (Obuća et al. 2022).

Importantly, this experiment different from our own in several key aspects: *i*) different suppliers of siRNA; *ii*) differences in cell types, i.e., HEK293 cells, whereas our experiment was conducted in HeLa cells; *iii*) shallower depth of sequencing, each sample yielding around 30 million uniquely mapped reads, compared to about 100 million reads in the case of our experiment; *iv*) shorter read length of 75 bps, compared to 126 bps. We note here that the last two aspects lead to a much lower coverage of the transcriptome (~5.5-fold). We processed this dataset with the same computational pipeline (see Methods), and quantified gene expression, as well as support for LSSs in order to learn whether downregulation of DHX38 leads to biased LSS activation similarly to what we observed in our experiment.

First, we found that quantification of DHX38 level indicates a slightly lower downregulation level of only ~1.5-fold, compared to about 2.2-fold that we observed in our experiment, which indicates lower effectiveness of the knockdown experiment and potential reduced downstream effects. Second, we found that the number of LSSs with split read support is much lower than in the case of our experiment, which is in agreement with the much more shallow depth of sequencing, given that transcripts using LSSs are much rarer species. Third, we found that activation of LSSs is not biased in samples where DHX38 was knocked down, in disagreement with our observations in HeLa cells.

**Effect of sequencing depth on detection of activated LSSs.**

The apparent disagreement between Obuća et al. and our study in terms of increased LSS activation upon DHX38 knockdown could have several explanations. First, different cell lines were used, HEK293 in the former and HeLa cells in the latter. Reduction in DHX38 mRNA levels were also different between the two experiments. More importantly, however, in the context of LSS activation is the aspect of lower transcriptome coverage in the RNA-seq experiment carried by Obuća et al. This is an important aspect because LSS-using transcripts are rare species due to their carrying in-frame STOP codons, and a high depth of sequencing is required to detect such transcripts. To test whether the depth of sequencing can significantly affect the ability of detecting increased LSS activation, we artificially lowered to depth of sequencing in our experiment to match that of Obuća et al. For this purpose, we randomly selected reads from the DHX38 and AA4 samples to match the average depth of sequencing in the Obuća et al. experiment, and then we retained for each read the 5'-most 75 bps to also match the read length. We then re-aligned these reads and quantified LSS usage using the same methodology as before. We obtained 30,445,793 uniquely mapped reads for the DHX38 sample, and 30,532,279 for the AA4 control sample. When we evaluated biased activation of LSS upon DHX38 knockdown compared to AA4 (non-specific siRNA), we found it to be not significant ($p=0.2$), despite the fact that using the entire set of reads (more than 100 million reads for DHX38) at their full length (126 bps) yielded a significant observation (Table S4). This finding indicates that a high depth of sequencing is critical in capturing the signal from the rare transcripts spliced at LSSs.
